# Immunological memory in a teleost fish: common carp IgM^+^ B cells differentiate into memory and plasma cells

**DOI:** 10.1101/2024.08.30.610468

**Authors:** Justin Tze Ho Chan, Amparo Picard-Sánchez, Neira Dedić, Jovana Majstorović, Alexander Rebl, Astrid Sibylle Holzer, Tomáš Korytář

## Abstract

From ancient cold-blooded fishes to mammals, all vertebrates are protected by adaptive immunity, and retain immunological memory. Although immunologists can demonstrate these phenomena in all fish, the responding cells remain elusive for lack of defining markers and tools to study them. Fundamentally, we posited that it is longevity that defines a memory cell like how antibody production defines a plasma cell. We infected the common carp with *Sphaerospora molnari*, a cnidarian parasite which causes seasonal outbreaks to which no vaccine is available. B cells proliferated and expressed gene signatures of differentiation. Despite a half-year gap between EdU labeling and sampling, B cells retained the thymidine analogue, suggesting that these are at least six-month-old resting memory cells stemming from proliferating precursors. Additionally, we identified a lymphoid organ-resident population expressing exceptional levels of IgM as plasma cells. Thus, teleost fish produce the lymphocytes key to vaccination success and long-term disease protection, and immunological memory is universal and universally demonstrable.

## Introduction

Over 500 million years ago, ancient cold-blooded fishes emerged along with adaptive immunity.^1^ Together with immunological memory,^2^ they form the basis of natural immunity, and vaccinology. These phenomena manifest as secretion of microbe-specific agglutinins, memory responses to antigen, accelerated rejection of allografts, and acquired protection from reinfection. They are observable and common to all fish including jawless fish,^3,4^ cartilaginous fish,^5–7^ and bony fish,^8,9^ with published reports dating back to over 80 years ago.^10^ Nowadays, we routinely observe adaptive immune and memory responses in a variety of fish species.^11–15^ Studying how these responses manifest promises solutions for preventing disease in fish. However, despite fish immunology coming to maturity, driven by comparative immunology and the importance of certain species to aquaculture, we still face difficulties identifying and studying the cells and mechanisms responsible for these phenomena.

It is especially difficult to identify homologous and functional molecules, cells and mechanisms in teleost fishes due to their rich diversity, evolutionary history and distance. For example, although the single-cell RNA sequencing of grass carp B cells identified plasma cells,^16^ there was a glaring absence of any population that could correspond to a memory B cell population. Fundamentally, it is phenomena such as longevity and antibody secretion that are observed time and time again rather than any specific phenotypic marker or mechanism. Therefore, it is from this angle that we studied the common carp B cell response, and searched for memory B and plasma cell analogues.

We leveraged our laboratory model infection of *Cyprinus carpio* with the myxozoan (Cnidaria) parasite *Sphaerospora molnari*.^17,18^ The myxozoans are ancient cnidarians that have co-evolved with and cycle between invertebrate and fish hosts. They cause economically important seasonal outbreaks in both wild and farmed fish stocks. In addition to the historical uncertainty over their phylogeny,^19–21^ the Myxozoa are also a scientific metazoan oddity due to adaptations such as one species losing the capacity for mitochondrial respiration.^22^ Generally, B cells respond strongly to myxozoan infections with proliferation and antibody production.^23,24^ Myxozoan infections potentially elicit polyreactive antibodies;^23^ they induce distinct transcriptional signatures that frequently include *il10* upregulation;^24–26^ infections are ultimately chronic and latent.^27^ These features of both the host response and the pathogenesis of myxozoan infections led us to question whether the B cell response elicits antibodies that mitigate disease and produces differentiated memory B and plasma cells that provide long-term protection. In other words, we investigated whether the B cell response is protective and productive.

In the present study, by removing B cells via immunosuppression,^28–30^ we provide evidence that B cells and antibodies limit the otherwise severe parasitemia and disease. Although we measured proliferation via incorporation of the thymidine analogue EdU and gene signatures that together indicate a productive B cell response, there are currently no specific phenotypic markers nor reagents to directly detect memory B cells. Thus, we relied on a single anti-carp IgM monoclonal antibody at our disposal. As described by Shibasaki et al. (2023) in a recent report of a germinal center analogue in fish, EdU labels cells engaged in an immune response and germinal center activity.^31^ It is through tracing the long-term retention of the thymidine analogue EdU that we directly detected resting memory B cells. A corticosteroid-resistant, enlarged, and head/anterior kidney lymphoid organ-resident B cell population expressing high levels of cytosolic IgM may represent plasma cells that constitutively secrete antibodies after resolution of the acute immune response. In this complex teleost host-myxozoan parasite interaction, humoral memory of a past infection, and protection are achievable.^32^ As our methodology is based on fundamental phenotypes of memory B and plasma cells rather than any homologous markers, it is universally applicable towards the study of these subpopulations in other species.

## Results

### 1.1 The B cell compartment expands and produces *S. molnari*-specific antibodies that limit parasitemia

To study the role played by B cells and the antibody response in *S. molnari* infection, we compared B cell-sufficient and B cell-depleted common carp throughout infection with the parasite. We previously revealed the most severe form of *S. molnari* infection via immunosuppression with the synthetic corticosteroid triamcinolone acetonide.^30^ Here, we demonstrate that immunosuppression completely depletes the peripheral IgM^+^ B cell population within 3 days of administration (Figure 1) and allows us to study the outcome of infection in the absence of B lymphocytes.

**Figure 1.**
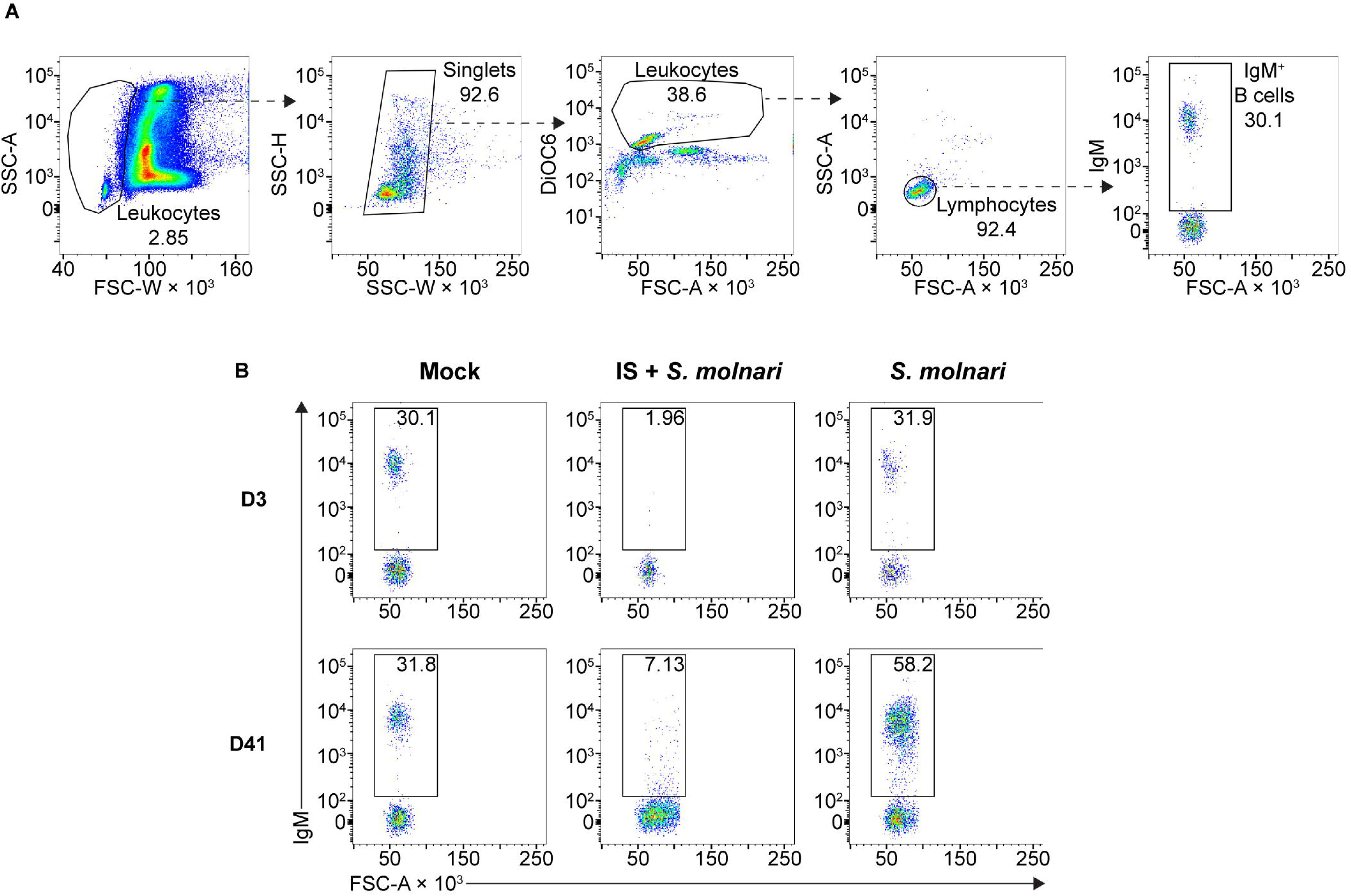
The circulating IgM^+^ B cell compartment of the common carp expands upon infection with the myxozoan *S. molnari* or is depleted upon immunosuppression. (A) A representative gating strategy for quantifying the IgM^+^ B cells in the whole blood of common carp. From the leftmost to the rightmost plots, dashed arrows each point out gated subpopulations (labeled with their proportions) being further analyzed in the plot to their right. The oval morphology of fish red blood cells gives them a heterogeneous scatter profile that overlaps with that of leukocytes. To exclude them, we first broadly gated on cells with low forward scatter width (FSC-W, x-axis), representing all leukocytes. Subsequently, doublets were excluded based on disproportionate side scatter (SSC) width (W) and height (H). Cells stained with the mitochondria- and endoplasmic reticulum-specific DiOC6 (y-axis) can distinguish active cells (leukocytes) from relatively inactive cells (red blood cells) with lower mitochondria or endoplasmic reticulum content. Among the leukocytes, the FSC^low^ SSC^low^ population represents lymphocytes which include IgM^+^ B cells based on staining with the monoclonal anti-carp IgM antibody WCI12 (mean fluorescence intensity, y-axis of the rightmost plot). Through this strategy, (B) we quantified the number of IgM^+^ B cells (gate and proportions indicated) throughout the experiment: from an early timepoint (day 3, top row) to a late timepoint (day 41, bottom row). Each column represents separate experimental groups either infected with 2,500,000 parasites (*S. molnari*), immunosuppressed and infected with an inoculum of 2,500,000 parasites (IS + *S. molnari*), or mock-infected (Mock). Representative plots are shown.

Fish were either not treated with the corticosteroid (immunosufficient) or immunosuppressed with it (IS). Under each of these two conditions, we further subdivided fish into three groups receiving one of three ten-fold diluted doses of DEAE cellulose-purified *S. molnari* BSs ranging from 2,500,000 to 25,000. Additionally, mock-infected fish were included as a control group.

Following infection, we measured changes in peripheral blood cellularity, focusing on expansion of the B cell compartment which is a hallmark of the infection and B cell activation (Figure 1 and 2A).^24^ Control fish that were mock-infected varied little throughout the experiment and little at early timepoints relative to the non-IS groups. When compared to this mock-infected group, the three IS groups were completely depleted of IgM^+^ B cells (Figure 1B, 2A, and S1). Lymphopoiesis only recovered and produced homeostatic B cell counts on day 41 post-infection in some groups and some individuals (Figure 2A). In contrast, the immunosufficient group had circulating B cells and this compartment began expanding as early as day 21 post-*S. molnari* infection. At this point, the number of peripheral blood IgM^+^ B cells increased gradually starting with the group that received the highest dose of 2,500,000 parasites followed by the group receiving ten times less, until IgM^+^ B cell cellularity was four-fold that of the control group on day 41 in the 6^th^ week of infection, the timepoint when we previously observed a peak of B cell expansion.^24^ Surprisingly, the blood B cell cellularity of the group infected with 25,000 BSs was not significantly different from the mock-infected group (Figure 2A and S1). The lowest dose of *S. molnari* BSs may be insufficient to establish an infection unlike the two higher doses. The dose-dependent expansion of the B cells may be a sign of B cell activation and an antigen-specific response.

**Figure 2.**
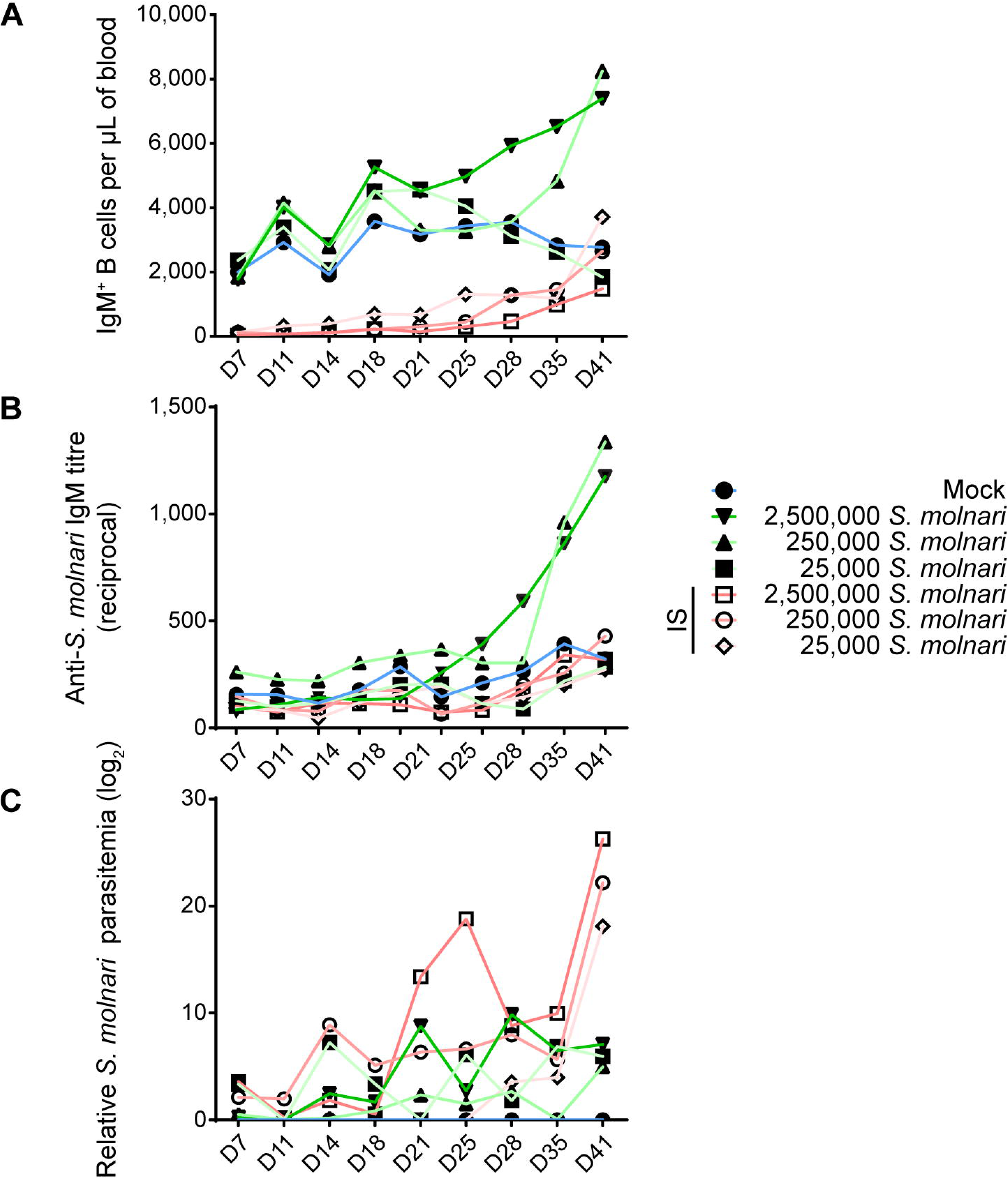
Infected common carp produce *S. molnari*-specific IgM which may limit parasitemia. By measuring (A) blood IgM^+^ B cell cellularity, (B) anti-*S. molnari* IgM antibody titres, and (C) parasitemia, we followed the progression of *S. molnari* infection in each of 6 groups of infected fish and one control group, represented by different line colors and symbols (legend on the right). We collected samples and took measurements at the indicated timepoints in days post-infection (x-axes). Filled or hollow symbols represent mean measurements from fish that were healthy or immunosuppressed (IS) respectively, at the time of infection; different shades of green or red as well as symbols represent different doses of *S. molnari* used in laboratory infection; the control mock-infected group is represented by blue lines. (A) Through the strategy depicted in Figure 1, we quantified the concentration of IgM^+^ B cells throughout the experiment. (B) The unit of measurement presented on the y-axis is reciprocal antibody titre, such that a higher number reflects a higher anti-*S. molnari* titre. (C) Parasitemia data was logarithmically transformed and presented in logarithmic units (base 2). n ≥ 5 biological replicates per timepoint per group. Please, refer to Figure S1 for the statistical analyses.

To measure the B cell response and the outcome of the infection, we quantified antibody titres and parasitemia throughout the experiment. Only baseline anti-*S. molnari* IgM antibodies were detected in the mock-infected and the IS fish (Figure 2B and S1). In immunocompetent fish, not only the number of B cells but also specific antibody titres were dose-dependent. Curiously, it was not only B cell numbers but also antigen-specific IgM titres that remained at baseline in the group that received the fewest parasites (Figure 2A and 2B), with a titre comparable to that of the mock-infected and all IS groups (Figure S1). We measured an increase in anti-*S. molnari* antibody titres as early as day 28 post-infection in the group receiving the highest dose of parasite (2,500,000 per fish), with the group receiving 10 times fewer parasites following shortly a week later. By the end of the experiment (41 days post-infection), specific antibody titres in these two groups were exponentially higher (less than 1:1000) than in any other group. Overall, these two groups had comparable anti-*S. molnari* IgM titres but significantly higher titres than any other group (Figure S1).

In IS fish, an exponential escalation of the infective dose (between 25,000 to 2,500,000) was matched by an exponential dose-dependent increase in parasitemia (Figure 2C and S1). The measurements of parasitemia in immunosufficient fish varied in kinetics (e.g., the timepoints at which parasitemia peaks). In these groups, the level of BSs in the blood of immunocompetent fish was ultimately not dose-independent unlike in IS groups. Parasitemia was not significantly different among these groups (Figure S1), and by day 41, all had over 2900-fold less parasitemia than any IS group.

Taken together, our infection model allows us to study the contribution of the B cells to the anti-parasitic response. Our data indicate that the immune system of the common carp rises to the challenge and suggests that it mounts a protective humoral/B cell response proportional to the parasite inoculum, with antibodies at least partly suppressing the parasite that would otherwise multiply unchallenged.

### 1.2 In response to *S. molnari* infection, IgM^+^ B cells proliferate predominantly in the lymphoid tissue, and express gene signatures of activation and differentiation

We shifted our attention to the splenic and head/anterior kidney lymphoid tissues where B cell responses are initiated. To characterize the B cell response, we measured proliferation and gene expression as indicators of activation and differentiation.

We detected significantly higher proportions of proliferating IgM^+^ B cells in all compartments studied (Figure 3). Proliferation uniformly peaked and was significantly different from corresponding control groups at week 5 post-infection in the blood, spleen, and head kidney (Figure 3B). This preceded the peak in both blood B cell cellularity and antibody titres by a week (Figure 2A and 2B). EdU incorporation and proliferation returned to baseline in all compartments two weeks later, indicating resolution of acute *S. molnari* infection. As for site specificity, the main compartment of proliferation at week 5 was the head kidney lymphoid organ where over 16% of IgM^+^ B cells had incorporated EdU versus about 7% in the blood and 8% in the spleen (Figure 3B). Proliferation being predominantly in the head kidney indicates that it may be a major site of B cell activation and differentiation in *S. molnari* infection.

**Figure 3.**
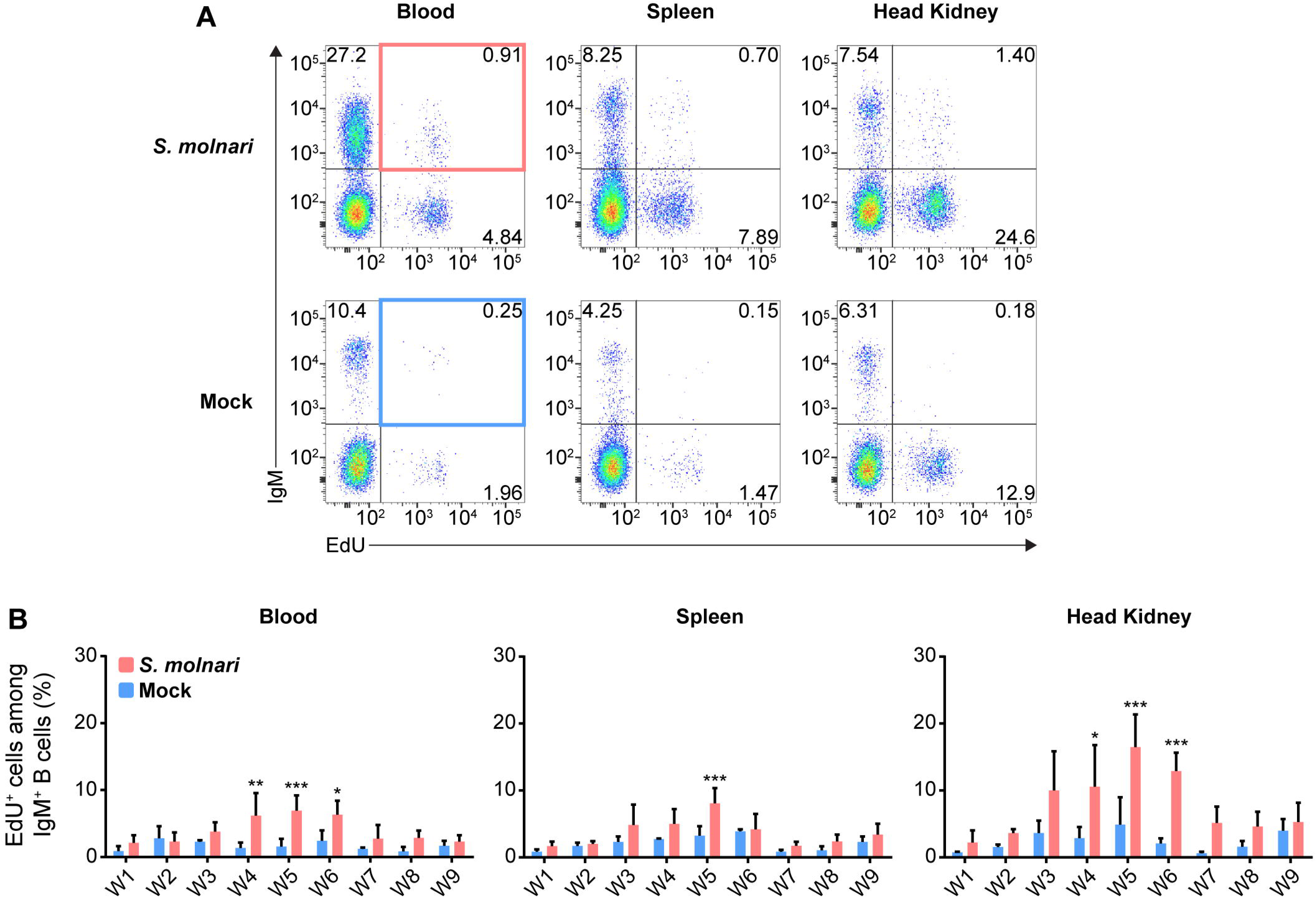
Common carp IgM^+^ B cells proliferate in the blood, the splenic lymphoid organ, and the head kidney lymphoid organ following *S. molnari* infection. (A) We exposed fish to the thymidine analogue (EdU) at different time points throughout *S. molnari* infection or in non-infected fish (Mock). As an indicator of proliferation, IgM^+^ B cells that incorporated EdU were identified by flow cytometry by adapting the gating strategy presented in Figure 1A to detect EdU^+^ cells in different tissue compartments. Representative plots from week 5 of the infection are presented here, organized by compartment (columns) and fish status (rows). The top two quadrants in every plot are the IgM^+^ B cell populations of interest. In the first column, the top right quadrants are marked by red or blue quadrilaterals which are the gates we used to quantify IgM^+^ EdU^+^ B cells from *S. molnari*-infected or control fish, respectively. (B) The proportion of IgM^+^ EdU^+^ B cells among total IgM^+^ B cells was quantified throughout the infection and summarized in bar graphs. Each bar extends to the mean with SD error bars. n = 3 or n = 4 biological replicates per timepoint for the Mock and *S. molnari* groups, respectively. ns (not significant); * *p* < 0.05; ** *p* < 0.01; *** *p* < 0.001.

To uncover and explain the changes that the B cells are undergoing in the head kidney throughout the infection, we profiled gene expression of magnetic-activated cell sorted (MACS-sorted) IgM^+^ cells (Figure 4A). We selected several predicted orthologues of mammalian and fish markers of B cell differentiation and/or survival (Figure 4B).^13,33–36^ Among the most differentially expressed genes (Figure 4B, top row, over 2-log change in relative gene expression), the significant downregulation of *membIgM* (membrane-bound or cell surface IgM), and upregulation of *secIgM* (secretory IgM), *xbp1*, and *tnfrsf13b* may together indicate differentiation of B cells into antibody-secreting and memory cells (Supplementary Figure 2). Specifically, the peak of *secIgM* and *xbp1* expression in week 5 post-infection corresponds with the timepoint when we began measuring an increase in anti-*S. molnari* IgM titres (Figure 2B) and may be indicative of B cell differentiation into antibody-secreting plasmablasts. The most differentially expressed gene was *tnfrsf13b* which was a significant 6 log units higher than the control group only at week 9 (Supplementary Figure 2). As a receptor in the BAFF/APRIL axis that promotes B cell survival and is also upregulated in rainbow trout by myxozoan infection,^37,38^ expression of *tnfrsf13b* (alias *taci*) may indicate the late differentiation of memory B cells. To date, no one has identified an orthologue of the human plasmablast marker *syndecan-1* (alias *cd138*) in teleost fish.^39^ We measured expression of the related *syndecan-3* but only observed that it was significantly downregulated at weeks 3 and 5. *pax5* and *cxcr5* expression were unchanged but the former trended toward downregulation at late timepoints. A two-way ANOVA determined that *secIgM*, *xbp1*, and *tnfrsf13b* expression were significantly higher in the head kidney than in the blood compartment.

**Figure 4.**
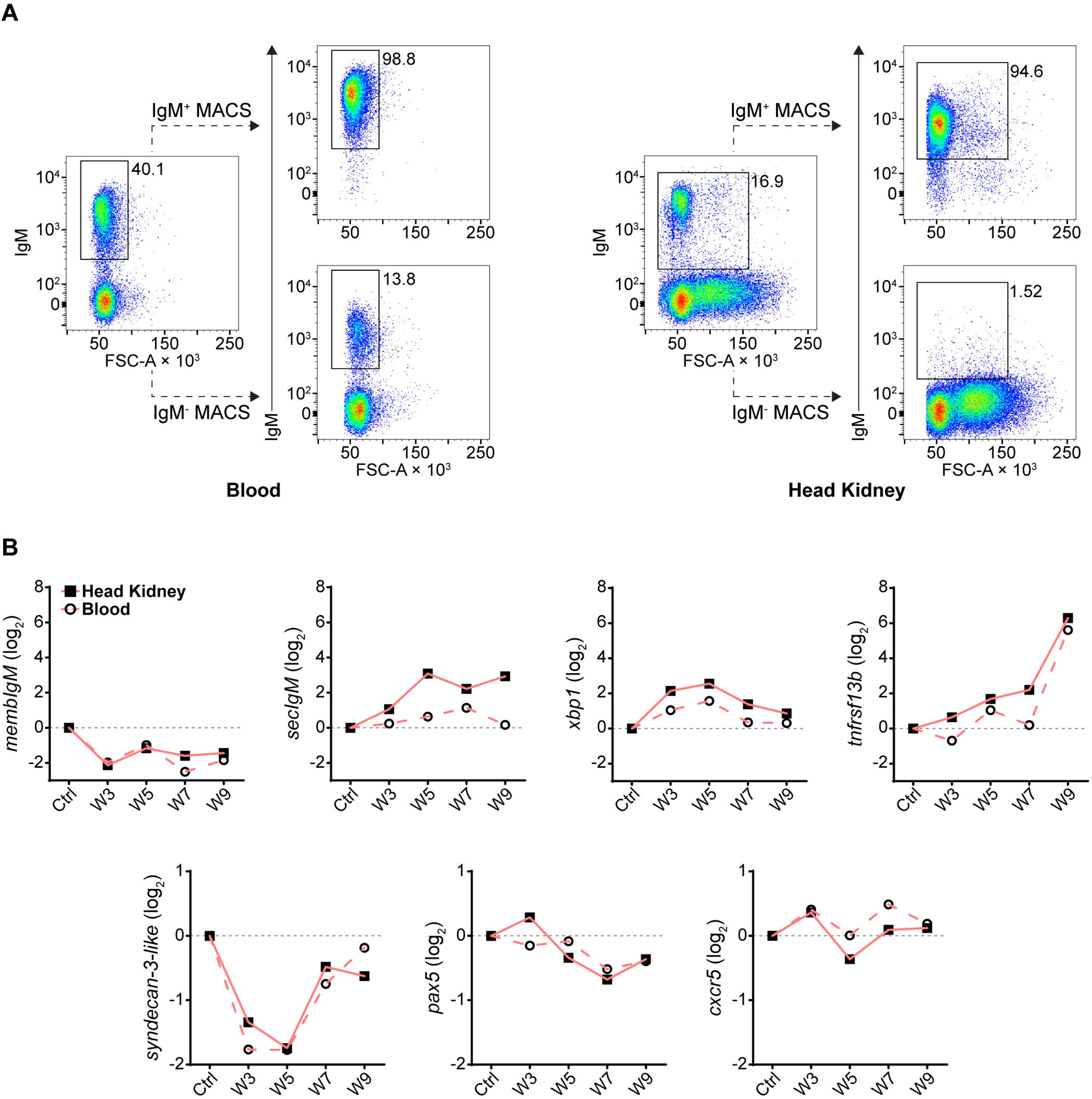
Following *S. molnari* infection, common carp IgM^+^ B cells express markers of B cell activation and differentiation in the blood and head kidney. (A) Head kidney or blood IgM^+^ B cells were MACS-sorted using the monoclonal anti-carp IgM antibody WCI12. Here are two representative MACS results for the blood (left) and the head kidney (right). In each plot, the proportion of IgM^+^ B cells is adjacent to the corresponding flow cytometry gate. The dashed lines point to the result of MACS: either the eluted IgM^+^ fraction (IgM^+^ MACS) or the unbound/washed negative fraction (IgM^-^ MACS). All plots depict the mean fluorescence intensity of WCI12 staining (y-axes) versus forward scatter area (FSC-A, x-axes). (B) RNA from MACS-enriched IgM^+^ B cells was used to quantify expression of select markers of activation and differentiation throughout *S. molnari* infection. Logarithmically transformed (base 2) relative expression (2^−ΔΔCT^ method) of each marker (y-axes titles) is presented in each plot and organized by highly or weakly differentially expressed genes: respectively, markers in the top row (over 2-log maximum change) or bottom row (under 2-log change). The control group (Ctrl) includes data collected from fish sampled prior to infection. We plotted the mean of relative expression of each group at each timepoint represented by either filled squares connected by lines (head kidney compartment) or hollow circles connected by dashed lines (blood compartment). n ≥ 4 biological replicates per group per timepoint. Please refer to Figure S2 for statistical analyses.

We further characterized these specimens via multiplex qPCR by profiling expression of B cell markers that were identified by Pan et al. (2023) in *Ctenopharyngodon idella* grass carp, another cyprinid species.^16^ Specifically, they are transcripts the authors identified via single-cell RNA sequencing of sorted head kidney IgM^+^ B cells. Here we present the expression levels of their potential *Cyprinus carpio* orthologues in our MACS-sorted IgM^+^ B cells (Figure 5). Expression of the 18 genes were hierarchically clustered within the three different compartments throughout the 10-week experiment for comparison between the blood, spleen, and head kidney compartments (Figure S3).

**Figure 5.**
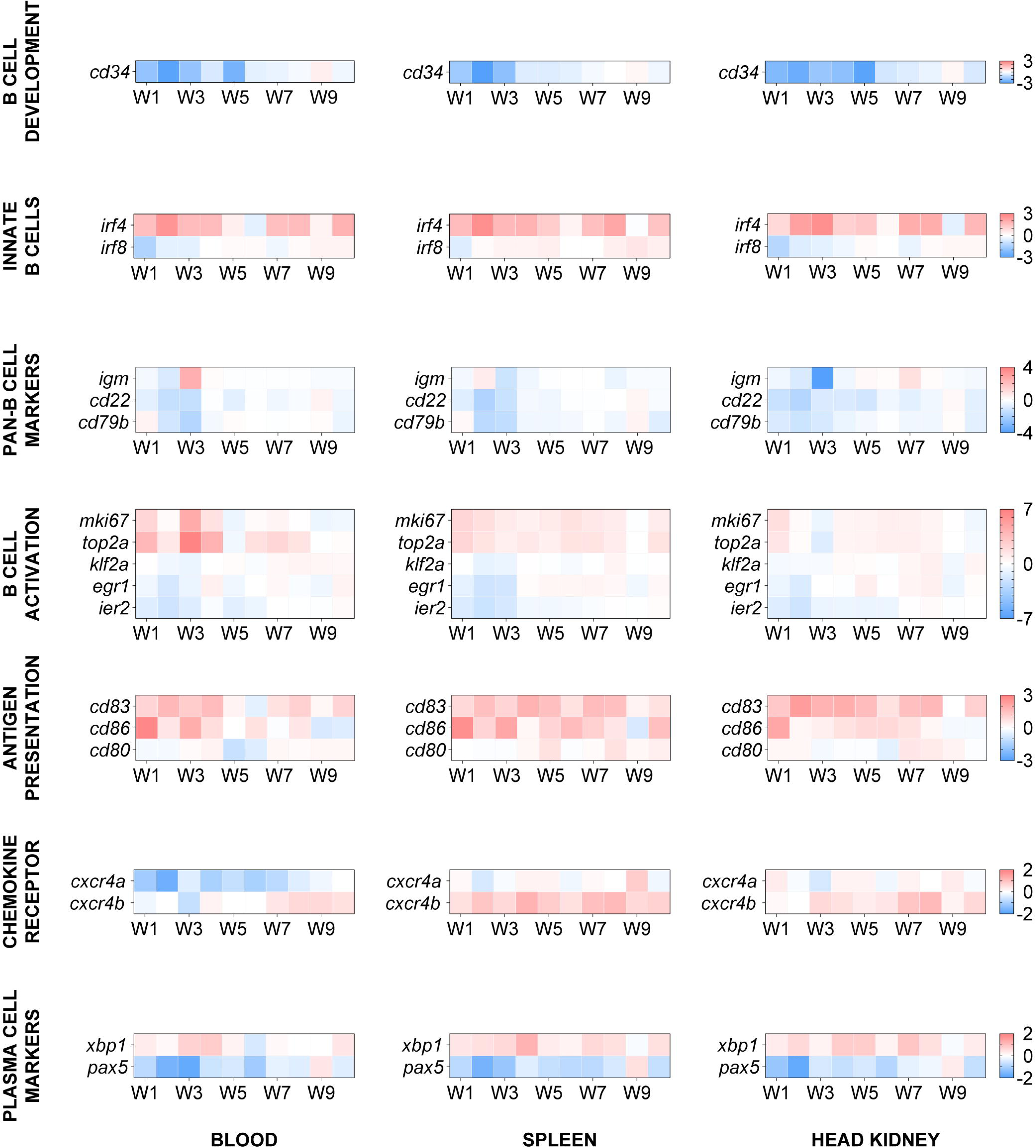
The transcription profile of cyprinid markers of developing, innate, activated/proliferating, antigen-presenting, and differentiated IgM^+^ B cell subpopulations. We profiled expression of markers defining distinct B cell subpopulations, i.e. genes initially identified in scrinia-seq analysis of grass carp IgM^+^ B cells.^16^ Here, genes were categorized (in separate rows of heat maps) based on the subpopulations they cluster, by developmental stage, and/or by activities such as activation and antigen presentation. The two-color gradient represents either overexpression or downregulation relative to the control group (not shown here). The legends for each gene category are on the rightmost side of each set of heat maps. Data in absolute number of copies of each gene were normalized to the number of copies detected in the control group and then logarithmically transformed (base 2) to better reflect both positive and negative fold changes. n ≥ 4 biological replicates per gene per timepoint. For statistical analyses, please refer to Figure S4.

The genes were categorized by their annotation and expected roles according to Pan et al.’s (2023) single-cell RNA sequencing data, and/or the cell population they help identify (Figure 5). The expression of *cd83*, *cxcr4b*, *egr1*, *klf2a*, and *ier2* were significantly different and the results of *post hoc* multiple comparisons tests are summarized in Figure S4. Notably, three of them are activation markers. *klf2a*, and *egr1* were downregulated at early timepoints (week 1 to 3); *egr1* was later upregulated between week 5 and 8 preferentially in the lymphoid organs while upregulation of *klf2a* was delayed and centered around week 8 (Figure S4). Among the co-stimulatory molecules, *cd83* was overexpressed in all compartments and throughout the experiment except the week 6 timepoint in blood IgM^+^ B cells unlike *cd86* which peaked at week 1 in all three compartments. *cxcr4b* was preferentially overexpressed at early timepoints in the lymphoid organs and may be the functional carp orthologue for retaining or directing select B cells to lymphoid tissues unlike *cxcr5* (Figure 4).

Since we are studying these markers at the population level and changes in IgM^+^ B cell subpopulations may still underlie statistically non-significant changes. c*d34* is a marker of mouse and human hematopoietic stem cells. It was downregulated in all compartments during the first 5 weeks of infection. Two markers of innate B cells, *irf4* and *irf8,* trended oppositely with the former highly overexpressed and the latter downregulated at early timepoints in all compartments. Among the pan-B cell markers, the most notable change was a 2.5-log upregulation and a 4-log downregulation of *igm* in the blood and head kidney, respectively, at the same week 3 timepoint. Similarly, the expression patterns of the activation markers *mki67* and *top2a* matched those of *igm* in the same cellular compartments at the same week 3 timepoint, suggesting that all three may be markers of the same IgM^+^ cell subpopulation. Finally, we again observed a reverse pattern of *xbp1* and *pax5* expression which is an indicator of an ongoing immune response and B cell differentiation (Figure 4 and 5).

In summary, considering that all these markers define distinct cyprinid IgM^+^ B cell subpopulations,^16^ our results suggest that there is: reduced hematopoiesis or lymphopoiesis; activation of an innate B cell population; antigen-presenting cell (APC) activity or costimulation; activation, proliferation, and differentiation of B cells; migration/retention of cells in lymphoid tissues. Together, these changes may be the origin of the humoral response against the parasite (Figure 2).

### 1.3 Memory B cells are produced throughout the acute response to *S. molnari* and persist as resting cells

Fish that survive a primary myxozoan infection presumably harbor memory lymphocytes that protect them from secondary infections. We gathered evidence that the initial immune response produces a greater breadth of B cells than we could previously appreciate with anti-carp IgM staining alone. Yet, we could not directly identify memory cells as there is so far no marker for let alone a defined subpopulation of teleost memory B cells. Furthermore, we must study these cells on the order or scale of months that are most relevant to immunological memory, vaccine success, and protection from natural seasonal (re-)infections. We revisited the EdU pulse labeling method, and searched for EdU^+^ IgM^+^ B cells months after infection which should theoretically reveal B cells that were i) activated upon encountering *S. molnari* (antigen), ii) proliferated and incorporated EdU as a result, and iii) were selected for and differentiated into long-lived quiescent cells that survived beyond the contraction and resolution phases of the primary immune response.

During acute *S. molnari* infection and proliferation of the B cells, we assigned fish to six labeling windows/groups receiving EdU during a specific week shortly after *S. molnari* infection. Approximately six months following infection and approximately five months following EdU injection, we nonetheless detected EdU^+^ IgM^+^ B cells in the blood, spleen, and head kidney compartments (Figures 6A and 6B). Astonishingly, this population was readily detectable in the lymphocyte gate, representing around 2% to 5% of all IgM^+^ B cells (Figures 6A and 6B) and potentially include memory cells. In contrast, a negligible amount of weakly EdU^+^ cells were detected among granulocytes from the same fish (Figure 6A). Compared to proliferation during the acute infection (Figure 3), the distribution of EdU^+^ cells was mainly in the blood and spleen rather than the head kidney, which matches the location of the resting B cells described by Ma et al. (2013);^13^ nor were they labeled during the 5^th^ week of acute infection, and peak proliferation (Figure 3 and 6C). The majority of EdU^+^ IgM^+^ cells originated from week 7 post-infection and at the time of measurement, represented 6% of all splenic IgM^+^ cells, and this number was about 4% and 3% in the peripheral blood and head kidney, respectively.

**Figure 6.**
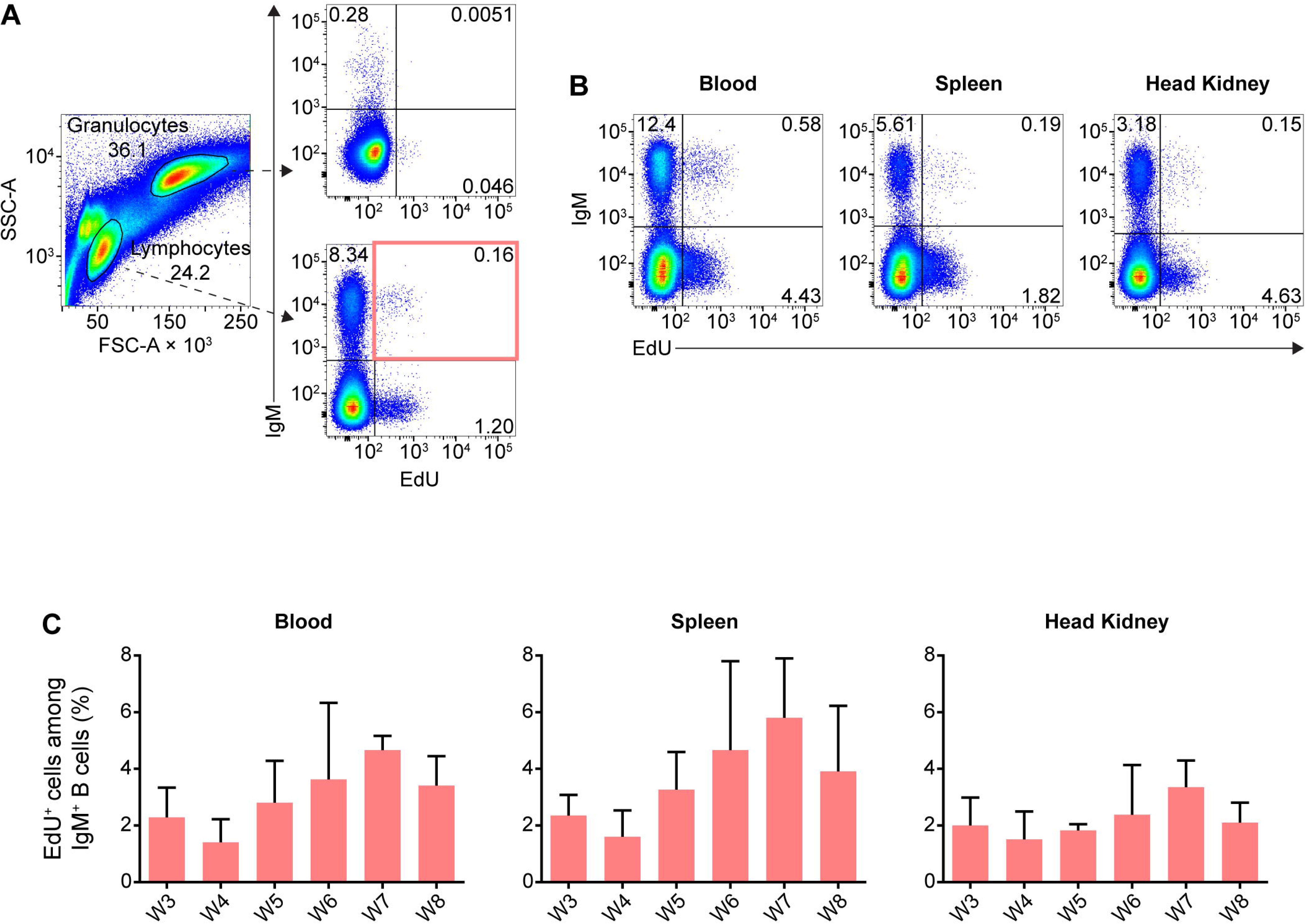
IgM^+^ B cells proliferating during acute *S. molnari* infection persist over five months later as resting cells. (A) In flow cytometry analysis of representative common carp head kidney leukocytes (left plot), distinct side scatter area (SSC-A, y-axis) and forward scatter area (FSC-A, x-axis) profiles help distinguish granulocytes (FSC-A^high^ SSC-A^high^) from lymphocytes (FSC-A^low^ SSC-A^low^). As indicated by dashed arrows, these two leukocyte populations were subsequently analyzed for/categorized by IgM expression (y-axis) and intensity of incorporated EdU (x-axis). Despite the months-long gap between EdU labeling and detection instead of next-day detection (Figure 3), EdU^+^ subpopulations were readily detectable among lymphocytes (bottom right plot), but rare among granulocytes (top right plot). (B) Additional examples of EdU^+^ IgM^+^ lymphocytes from the blood, spleen, and head kidney are presented. Proportions of each quadrant/subpopulation are noted on plots. In (A) we highlighted the main EdU^+^ IgM^+^ population of interest via a red rectangle and the proportion of this subpopulation relative to total IgM^+^ lymphocytes is summarized in bar graphs in (C). These graphs are organized along the x-axes by separate bars (mean + SD), each representing a labeling window: the week post-*S. molnari* infection in which each group was injected with EdU. Representative data presented from the week 4 and 7 labeling windows in (A) and (B), respectively. n ≥ 3 biological replicates.

In other words, out of all the proliferating and EdU-incorporating cells during acute infection, a subset survived, having not been turned over, not proliferated further, and not diluted EdU below the threshold of detection. This B cell population may be partly composed of the fish equivalent of the memory B cell, a non-dividing recirculating B cell subset awaiting reactivation.

### 1.4 A head kidney-exclusive B cell subpopulation with high levels of intracellular IgM may represent plasma cells

Plasma cells and their antibody effector molecules make up another pillar or wall of humoral memory. We were able to detect anti-*S. molnari* antibodies eight months post-infection, long after resolution of parasite infection and without any reinfection or immunization (Figure 7A). With these antibodies being presumably produced by plasma cells, we devised methods to detect this B cell subset based on the exceptional amounts of Ig they produce, their lower levels of membrane IgM (membIgM) or B cell receptor (BCR),^35^ their size, their compartmentalization in the head kidney,^13^ and resistance to hormonal stress.^40^

**Figure 7.**
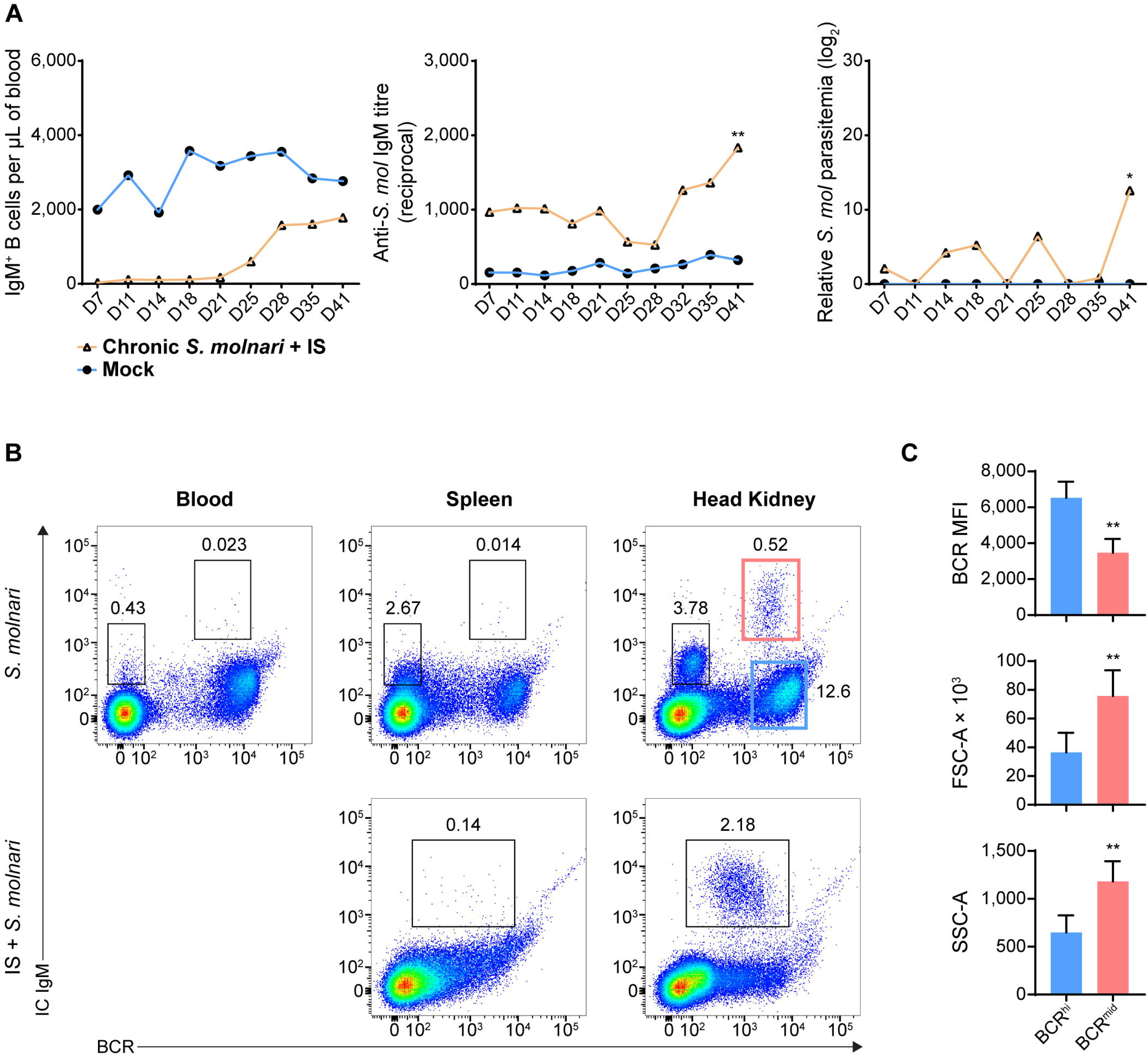
A population of head kidney-specific large B cells with exceptionally high levels of intracellular IgM and resistance to immunosuppression may represent plasma cells of the common carp. (A) We studied a cohort of immunocompetent carp infected eight months prior which have not achieved sterile immunity (labeled ‘Chronic *S. molnari*’). Unlike fish infected for the first time (Figure 1, the ‘Mock’ control group was sampled simultaneously as the ‘Chronic *S. molnari’* group), these carp constitutively produced anti-*S. molnari* IgM (middle plot) which are conventionally produced by plasma cells. As we hypothesized that plasma cells are resistant to immunosuppression, the ‘Chronic *S. molnari*’ cohort of fish was treated with triamcinolone acetonide (IS) which did not decrease specific antibody titres throughout the experiment despite completely depleting circulating IgM^+^ B cells (left plot). Nonetheless, parasitemia (right plot) was measurable in these fish. Orange line and triangle symbols represent the *‘*Chronic *S. molnari* + IS’ group whereas the blue line and circles represent an uninfected control cohort mock-infected (Mock). (B) We identified a population of putative plasma cells via costaining for membrane/cell surface IgM (the BCR, x-axes) and intracellular IgM (IC IgM, y-axes) lymphocytes. This IC IgM^high^ population was detectable exclusively in the head kidney (rightmost column). Despite near-complete depletion of IgM^+^ B cells in the blood (Figure 1B and 2A), and head kidney, triamcinolone acetonide treatment (IS) enriched this IC IgM^high^ population in the head kidney (bottom right plot). (C) Each bar indicates the mean of either the mean fluorescence intensity (MFI) of membrane IgM staining, forward scatter area (FSC-A), or side scatter area (SSC-A) + SD. For these metrics, statistically significant differences between the BCR^high^ and BCR^mid^ populations, respectively outlined in blue and red rectangles in (B), were determined by the Wilcoxon signed rank test with the two populations as matched pairs in n = 8 biological replicates. ** *p* < 0.01.

Thus, we costained cells in a step-wise manner to detect the extracellular BCR and intracellular IgM (IC IgM) using a single monoclonal anti-carp IgM antibody. In the blood and spleen, the B cells stained minimally for IC IgM with a slight upward shift of the BCR^high^ population of cells (Figure 7B, top row). This did not reveal any new population among the cells we have always detected (Figure 1). However, the head kidney revealed two new B cell populations that were previously undetectable through BCR staining alone (Figure 7B, top row): a BCR^-^ IC IgM^mid^ population and a BCR^mid^ IC IgM^high^ population. The former is relatively abundant (representing around 3% of all leukocytes) while the latter is exceedingly rare (representing around 0.5% of all leukocytes). Overall, based on their phenotype and tissue-specificity, we hypothesized that one of these populations represents plasma cells.

To test these cells’ resistance to stress, we immunosuppressed fish that had previously been infected with *S. molnari*. As the corticosteroid-induced immunosuppression targets lymphopoiesis,^41^ dividing cells, and short-lived cells, we hypothesized that plasma cells would be resistant and would not be depleted unlike their naïve B cell counterparts. This treatment eliminates virtually all IgM^+^ B cells from the blood (Figure 1B and 2A). In the spleen and head kidney, this approach depleted the most abundant B cell populations including the BCR^-^ B cell population (Figure 7B, bottom row). In contrast, the BCR^mid^ IC IgM^high^ population became readily detectable in the head kidney (over 2% of all leukocytes in the example). We also hypothesized that they would continue producing antibodies despite the immunosuppression and were responsible for keeping the parasite latent long-term. Thus, we measured antibody titres and parasitemia in the weeks following immunosuppression (Figure 7A). Anti-*S. molnari* IgM antibody titres doubled and coincided with the reemergence of the parasite and an over 10-log increase in parasitemia at the final timepoint of 41 days post-immunosuppression.

Together, our data indicate that the initial B cell response (Figure 1 and 2) elicits constitutive antigen-specific antibody production. The source of these antibodies may be the corticosteroid-resistant, head kidney-resident, IC IgM^high^ B cell population, which are significantly larger and denser than their counterparts with high BCR expression (Figure 7C). These cells and antibodies may be one component of constitutive defense keeping *S. molnari* latent and/or protecting the host from future infections.

## Discussion

Overall, our data indicate that B cell activation follows a similar course to that in ‘higher’ vertebrates and produces specialized cellular subsets with memory and (constitutive) antibody-secreting phenotypes, together forming ‘two walls of protection’.^42^ Our results indicate that the initial B cell response is protective and prevents the worst form of disease caused by *S. molnari* (the only two deaths recorded in this experiment were from the IS groups).

### 2.1 The B cell response, memory B and plasma cells in a teleost fish

In mammals, antigen activates B cells and they differentiate in lymphoid tissues. This process can be divided into at least two phases: an initial phase that produces the short-lived plasmablasts and some memory B cells, whereas a second phase produces memory B and long-lived plasma cells that have affinity-matured and have been positively selected for in germinal center reactions.

The B cell response we observed was dose-dependent, and antigen-specific. In comparison to the response of rainbow trout to the model hapten-protein antigen TNP-KLH, anti-*S. molnari* antibody production was accelerated, and titres were exponentially higher at the same timepoint compared to the response of rainbow trout against TNP-KLH.^12^ The initial surge in antibody production at the 5-week timepoint (Figure 2B) is likely the product of short-lived plasma cells or plasmablasts and matches the timing of the significant increase in *igm* transcripts we previously observed in the same model and in this study (Figures 4 and 5).^24^

We measured a peak of proliferation at week 5 post-infection and detected a significant proportion of EdU^+^ IgM^+^ B cells in both the head kidney and spleen. These cells likely include those described in a recent report by Shibasaki et al. (2023) of a germinal center analogue in the rainbow trout spleen.^31^ By detecting highly proliferating B and T cells adjacent to melanomacrophages, Shibasaki et al. convincingly identified sites that support clonal expansion, antigen receptor affinity maturation, and clonal selection. One expected output is antibody-secreting cells which circulate and contribute to the sharp rise in B cell cellularity and EdU^+^ B cells at week 5 and 6 in the blood even though it is not a site of proliferation (Figures 2A and 3). The short-term distribution of antibody-secreting cells throughout the periphery was also observed by Davidson et al. in *Limanda limanda* following immunization with human gamma globulin.^11^ The precise identity of the circulating B cells can be confirmed with a strategy like the one we used to detect the kidney-resident IC IgM^high^ cells. However, they are only expected to be short-lived and provide short-term protection.

Regardless of which phenotypic markers they express, regardless of species-to-species variations, the purest and most universal definition of what is a memory B cell is their longevity. We traced the proliferating cells of the primary response to determine if they persist long after resolution of the acute response. We hypothesize that the EdU^+^ cells we detected months after infection/labeling include clones that avoided caspase-mediated apoptosis of the melanomacrophage centers.^31^ The upregulation of *secIgM*, *xbp1*, *tnfrsf13b,* and downregulation of *membIgM*, and *pax5* may also be byproducts of ongoing selection and differentiation. We detected memory B cells in every compartment, and they formed at every timepoint studied. The memory B cells detected in the spleen and blood matches where Ma et al. (2013) observed them in rainbow trout.^13^ These cells are likely recirculating between the two compartments, explaining why their proportions are very similar in the two compartments. In contrast, the head kidney of carp and other bony fish species appears to be a niche for plasma cells that do not recirculate. These cells likely reside in the head kidney which is a survival niche for plasma cells of other bony fish species as well.^13,14,16,43^

The emergence of memory cells and the peak of their formation in week 7 (Figure 6C) was relatively late and did not correlate with the peak of B cell proliferation in week 5 (Figures 2B and 3). Considering the timing and selection mechanisms ongoing in lymphoid tissues, it is likely that early memory B cells express low affinity BCRs. In contrast, the late-stage memory cells that emerged in week 7 endured activation-induced cytidine deaminase-mediated antigen receptor diversification, express affinity-matured BCRs, and have a survival/fitness advantage as successful progenitors of melanomacrophage center reactions.

### 2.2 Humoral memory and the implications for vaccination and natural immunity

In humans, there is evidence that exposure to a pathogen is sufficient to protect an individual for a lifetime: in a 2008 study, survivors of the 1918 H1N1 influenza pandemic continued to harbor memory cells producing strain-specific anti-hemagglutinin neutralizing antibodies, nearly 90 years after exposure to the virus.^44^ If fish B cells behave in much the same way, there are major implications for vaccination and natural acquired immunity against pathogens. A similar study in mice detected BrdU retention in memory B and plasma cell populations at the 8-week timepoint.^45^ Here, we continued to detect EdU^+^, IgM^+^ B cells well after 6 months post-infection and EdU pulse labeling. This progress and this milestone are consequential because we need to know how long memory is maintained post-vaccination or whether fish having already survived an initial infection will be immune during next year’s outbreak. Understanding how memory is generated and maintained will help us move away from purely empirical vaccines and methods, and towards conferring acquired immunity.

Existing models propose that memory B cells could be maintained by continuous antigen exposure while others propose that their longevity may be intrinsic to ‘tonic’ BCR signaling. Although the former suggests that the immune system retains antigen from a primary response, we may also artificially re-expose B cells to provide survival signals via booster immunizations. The germinal center-like phenomenon likely applies to immunization as well: following TNP-keyhole limpet hemocyanin immunization, Ye et al. demonstrated that the average affinity of the rainbow trout B cell population matures over time.^12^ Theoretically, the reactive humoral memory provided by memory B cells is not completely redundant with that provided by constitutive memory (antibodies). Considering the discovery of a germinal center analogue in fish, the fish memory B cells we detected may potentially also re-enter melanomacrophage centers to further diversify their antibody repertoire and/or differentiate into (short-lived) plasma cells, providing a ‘faster’ and ‘stronger’ secondary immune response.

Finally, there is the issue of what our findings mean for natural infection or immunity. We do not know if the survival of memory cells in laboratory settings is reproducible in natural settings with all its environmental stressors and the potential for sequential infections and coinfections. Interestingly, Zwollo (2012) hypothesizes that one stressor is the immunosuppressive and B cell-depleting cortisol produced by spawning fish.^40^ However, the long-lived plasma cells are exceptionally resistant to the corticosteroid activity and continue providing humoral protection. In this ‘immunological imprinting’ scenario, long-lived plasma cells were generated in juvenile fish against pathogens in their rearing site. As they return to these sites as spawning adults, the long-lived plasma cells are undeterred by hormonal changes, and continue producing antibodies in anticipation of familiar pathogens. Another study demonstrated that rainbow trout plasma cells are resistant to hydroxyurea.^14^ The implications are that these long-lived resistant plasma cells may be key to interseasonal protection of fish in natural settings. In our study, we mimicked the effects of cortisol in the previously *S. molnari*-infected fish. Yet, antibody titres remained stable (Figure 7A) without *de novo* B cell lymphopoiesis, without circulating naïve B cells, and without *de novo* antibody production, despite the half-life of fish IgM being reportedly mere days.^46^ Thus, it suggests that anti-*S. molnari* plasma cells would survive such a stressor and that their antibody production is not affected (Figure 7B). It may be one underlying mechanism protecting immune adult fish versus naïve juvenile fish. As the plasma cells are presumably terminally differentiated without capacity for reactivation, the rise in antibody titres after weeks of stability in the experiment (Figure 7A), may indicate that memory B cells were also hormone-resistant and were reactivated by reemerging parasites.

### 2.3 Potential lessons from the *S. molnari* infection model

Chronic infections pose the greatest challenges even in well-studied species. For example, despite decades of concentrated research efforts, there is still no vaccine available against pathogens causing chronic diseases such as human immunodeficiency virus and malaria. Interestingly, in malaria infection, despite robust production of plasmablasts and anti-*Plasmodium* antibodies, this was ultimately metabolically taxing and impaired production of memory B and plasma cells.^47^ Our data suggests that we can rule out this possibility to explain chronic *S. molnari* infection because we measured a persistent constitutive anti-*S. molnari* response. The immune antisera from infected fish can lyse the parasite (manuscript in preparation) and likely keeps *S. molnari* latent.

In natural settings, could immunosuppression (e.g., via cortisol) be a trigger for parasite activation, spore formation, and perpetuating the life cycle? Sterilizing immunity to myxozoan infections is rare and fish become carriers, suggesting that the parasite can evade the immune system, potentially a general feature of myxozoans.^48^ Despite continued secretion of anti-*S. molnari* antibodies, humoral memory could not provide sterilizing immunity nor keep the parasite latent eight months later (Figure 7A). It is possible that plasma cells expressing antibodies specific to an original ‘founder’ variant could no longer recognize an escape variant of the parasite (manuscript in preparation). Thus, if fish return to the same familiar and reliable spawning location yearly and it is a site where the invertebrate host is found as well, myxozoan parasite escape and sporulation may be side effects of the ‘immunological imprinting hypothesis’ and hormonal immunosuppression. In this model, interseasonal carriers such as the common carp or wild trout may spawn and produce invertebrate-infective spores in spring.

Infected invertebrates may produce the fish-infective spores, completing the parasite life cycle and causing the annual outbreaks of myxozoan infections.

In summary, we adapted and devised methods that may be applicable to other vertebrate species in which adaptive immunity and immunological memory can be demonstrated because it relies on a common phenomenon, not a common mechanism or marker. Perhaps what holds true for all vertebrates and the only all-encompassing definition is that memory lymphocytes are antigen-experienced, born of an immune response, and persist after resolution of that response. Immunologists have observed immunological memory for over a century and achieving humoral memory will help us meet the needs of immunologists, parasitologists, and fish husbandry.

## Materials and methods

### 3.1 Key resources table

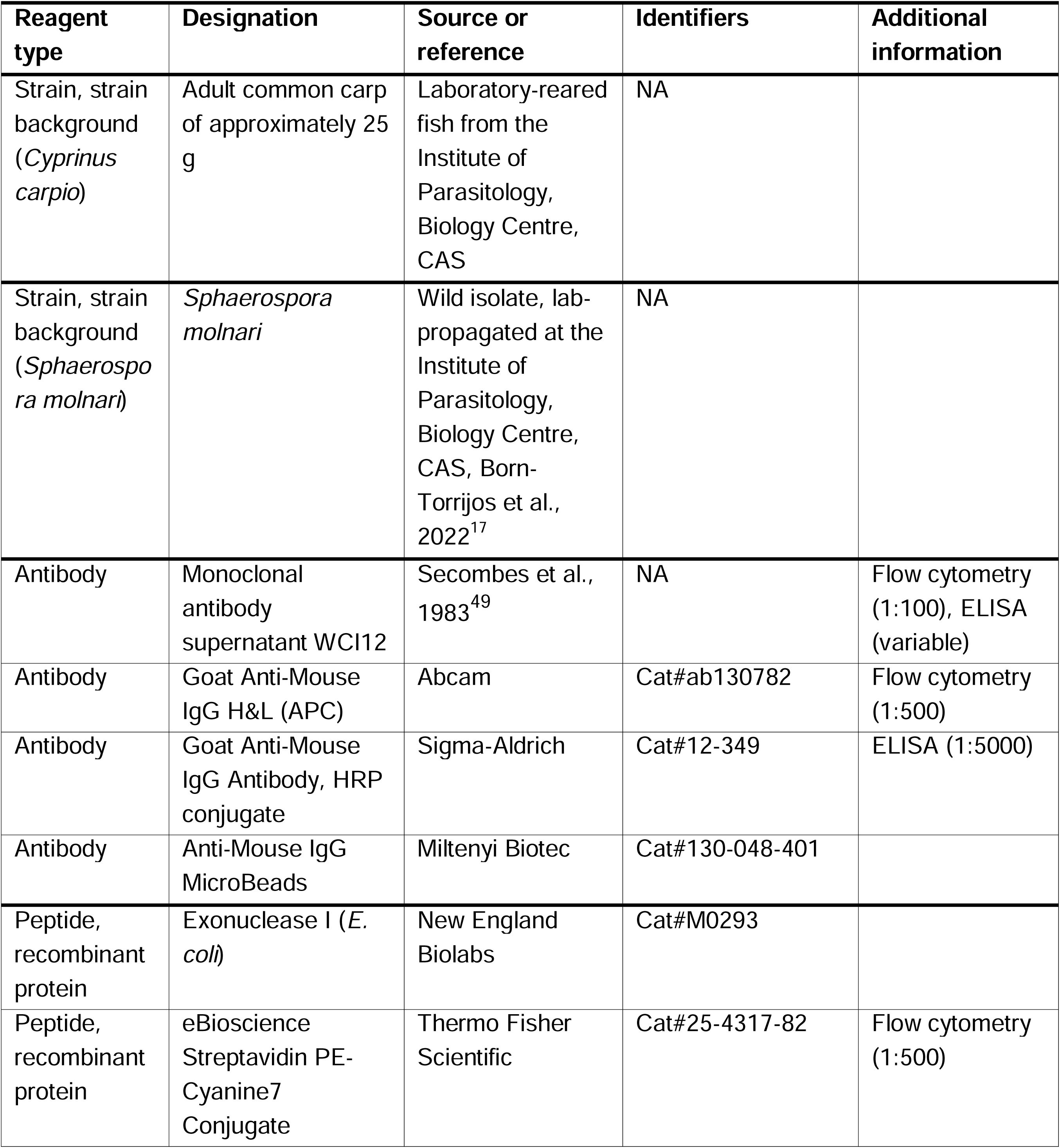
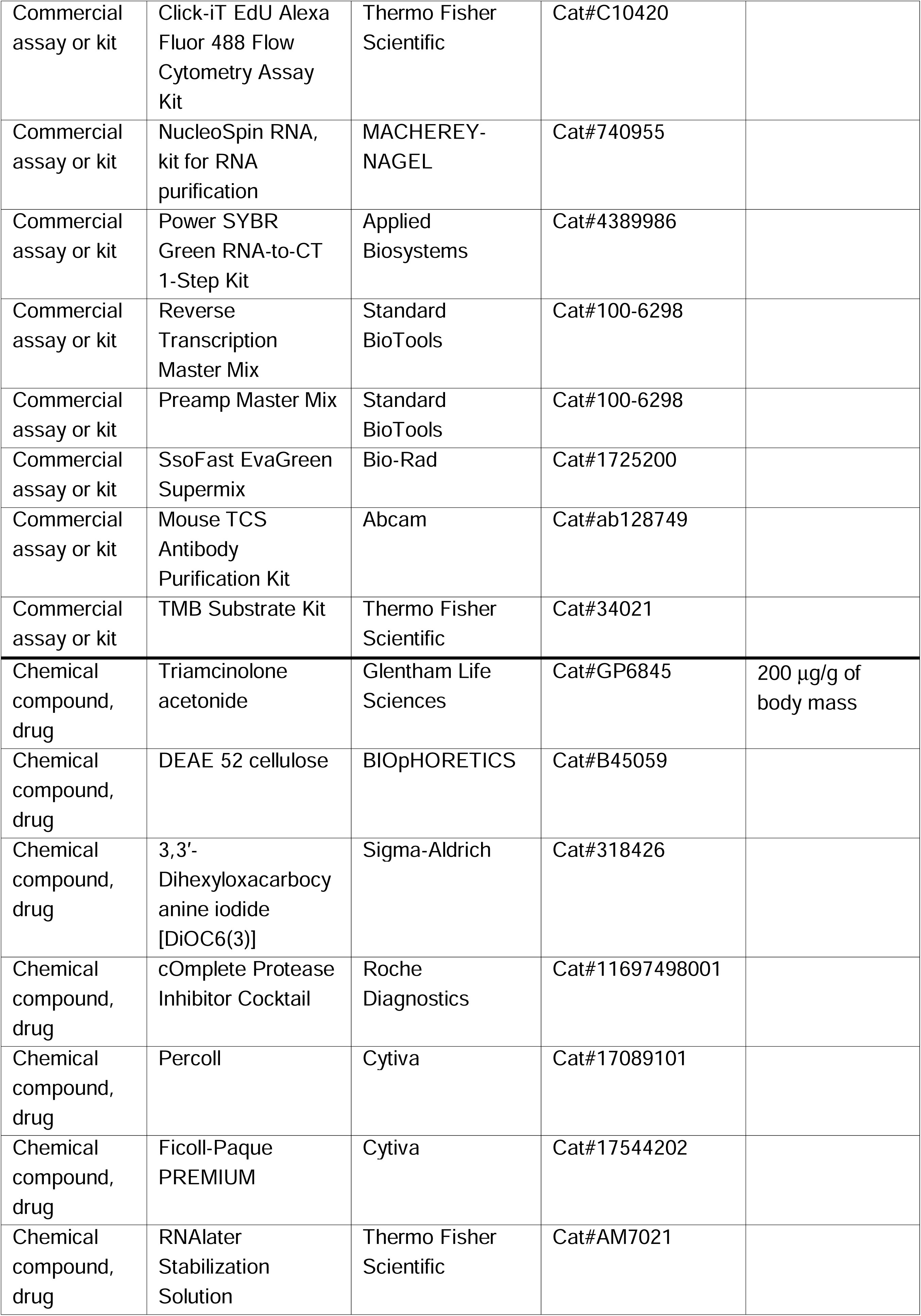
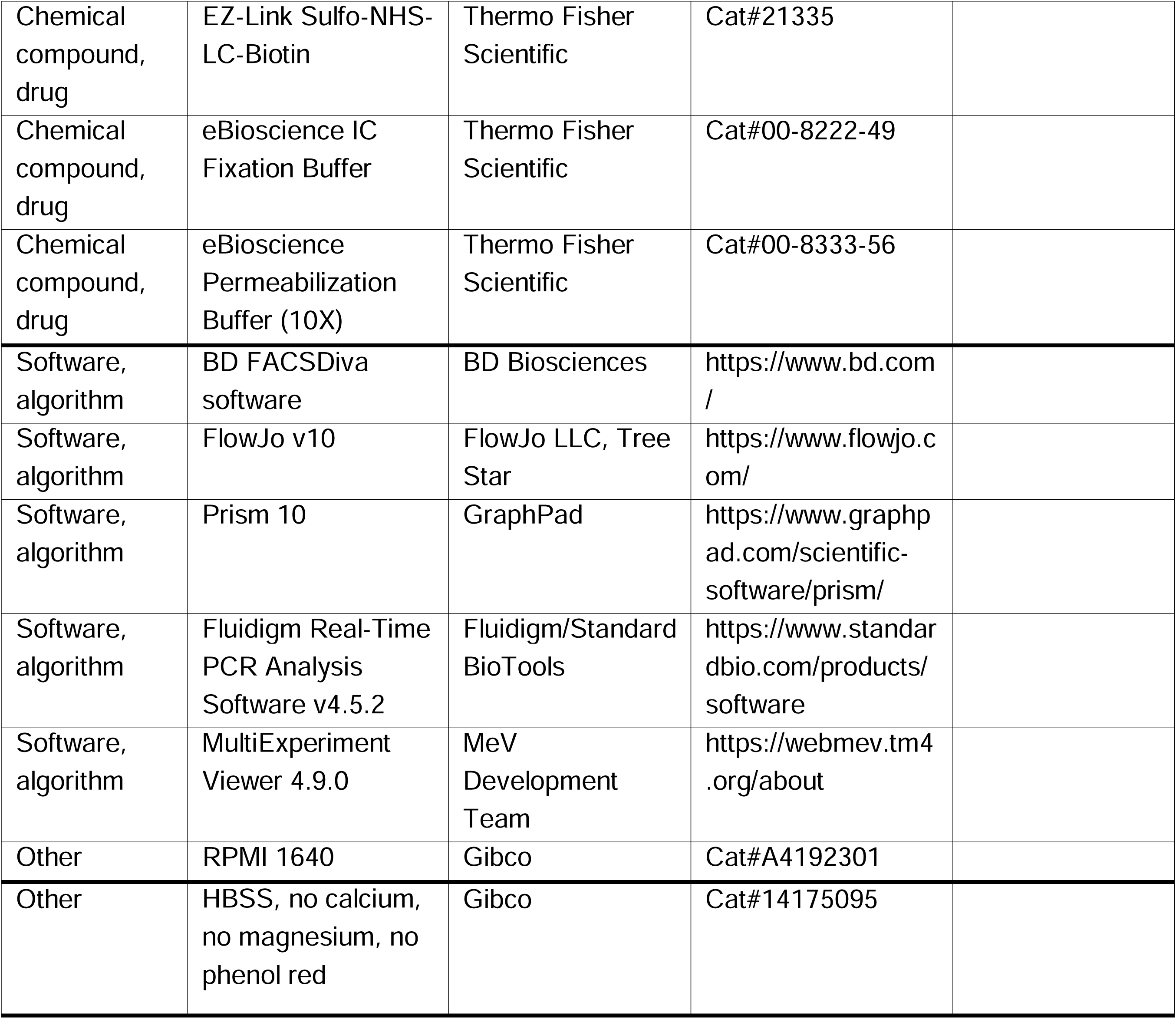

### 3.2 Experimental model details

Animal procedures were performed in accordance with Czech legislation (section 29 of Act No. 246/1992 Coll. on Protection of animals against cruelty, as amended by Act No. 77/2004 Coll.). Animal handling complied with the relevant European guidelines on animal welfare (Directive 2010/63/EU on the protection of animals used for scientific purposes) and the recommendations of the Federation of Laboratory Animal Science Associations. The animal experiments were approved by the Ministry of Education. Approval ID: MSMT-4186/2018-2. Our study is reported in accordance with ARRIVE guidelines (https://arriveguidelines.org).

We reared specific pathogen-free (SPF) common carp (*Cyprinus carpio*) from peroxide-treated fertilized eggs (700 mg/L for 15 min) in an experimental recirculating system in the animal facility of the Institute of Parasitology, Biology Centre CAS. During the experiment, fish with an average mass of approximately 25 g were selected and fed twice a day with a commercial carp diet (Skretting) at a daily rate of 1.5% of their body mass. The sex of experimental fish was not considered in our study. The fish were implanted with passive integrated transponders to track the experimental group they belonged to. We housed fish randomly and therefore fish from different experimental groups shared the same tanks. However, all tanks shared the same water which was UV-irradiated, ozonized water at 21 ± 1 °C, with water quality (oxygen, pH, ammonia, nitrite and nitrates) monitored daily using probes and titration tests. Ammonia levels never surpassed 0.02 mg/L.

### 3.3 Experimental infections and fish

For all infections, we collected *S. molnari* parasite from the fish host and purified the blood-stage parasite cells (BSs) via DEAE cellulose (BIOpHORETICS, Sparks, NV, USA) isolation as described previously.^17^

For the trial measuring B cell cellularity, antibody titres, and parasitemia, fish were divided into eight groups of six fish each for the initial kinetic measurements of peripheral blood cellularity, antibody titres, and parasitemia: groups 1-3 were immunocompetent/immunosufficient and received one of three ten-fold diluted doses of parasite ranging from 2,500,000 to 25,000; groups 4-6 were also given one of the three doses of parasite as described for groups 1-3 but these fish were immunosuppressed with triamcinolone acetonide (Glentham Life Sciences, Corsam, UK) at a dose of 200 µg/g of body mass the day before intraperitoneal injection of the parasite; ^17^ finally, one group was mock-infected via injection of Roswell Park Memorial Institute 1640 medium (RPMI 1640, Life Technologies Limited, Paisley, UK) that also served as the medium for the parasite.

For the downstream EdU labeling assay, we infected two additional groups of fish with 50,000 BSs per fish, and mock-infected one more group. One group was mock-infected with RPMI 1640 (n = 27) and served as a control for the second group infected with *S. molnari* (n = 36). The final third group was divided into six sub-groups (n = 4 each) for the purpose of memory B cell detection.

To measure differential gene expression, 50 fish were infected with one million BSs each, five fish for each of ten sampling timepoints, sampled at one-week intervals for the ten-week duration of the experiment. An additional five non-infected fish were dissected and served as the control group.

A final independent cohort was infected with 50,000 BSs per fish before immunosuppression rather than immunosuppressed first. We subjected this group to a 223-day gap between infection and treatment with triamcinolone acetonide (as described above) to study immunological memory.

### 3.4 Measuring peripheral blood B cell cellularity

We collected minimal quantities (around 50 to 100 μL) of blood from experimental fish at a frequency of twice weekly throughout the 6-week experiment. The blood was collected with 30-gauge needles into heparinized syringes (via rinsing of the needle and syringe with 200-300 μL of 5,000 IU/mL heparin sodium before use). In preparation for flow cytometry, 4 μL of the whole blood specimens were aliquoted, the blood was resuspended in 200 μL of RPMI 1640 before 100 μL was discarded. Preparation of specimens for flow cytometry was in the 96-well plate format. We centrifuged the cells at 500 *g* for 3 min and washed them once more to dilute and remove any serum (all washes and stains were in the absence of antibiotics or bovine serum). In parallel, 2 μL of the whole blood was collected in TNES-urea to measure parasitemia. The remainder of the heparinized blood was centrifuged at 500 *g* for 5 min. We collected and froze the plasma at -20 °C until necessary for downstream applications.

For flow cytometry, the cells were then stained with previously titrated SP2/0 hybridoma supernatant containing the monoclonal anti-carp IgM antibody (clone WCI12)^49^ diluted 1:100 in RPMI 1640. We incubated cells for 20 min in ice before subjecting cells to two washes to remove excess antibodies. 1:500 goat anti-mouse IgG1-APC (Abcam, Cambridge, UK) was then applied as a secondary detection reagent, also diluted in RPMI 1640. Incubation was for 20 min in ice before we spun cells down, and added 3,3’-dihexyloxacarbocyanine iodide [DiOC6(3)] (Life Technologies, Carlsbad, CA, USA) diluted to 5 μg/mL in RPMI 1640. We used this lipophilic dye as an alternative to any density centrifugation step, to distinguish the relatively metabolically active and endoplasmic reticulum- or mitochondria-rich leukocytes from the relatively inactive red blood cells. All stains were performed in a volume of 100 μL. A final centrifugation took place before we resuspended cells in a final volume of 200 μL of RPMI 1640. The cells were measured on the BD FACSCanto II (BD Biosciences, Prague, Czech Republic) and data was recorded for 20 s at a medium flow rate (60 μL/min) for all samples, allowing us to quantify both proportion and absolute numbers of B cells in the original 2 μL of whole blood.

We used a two-laser configuration of the BD FACSCanto II: equipped with a 488-nm blue laser and a 633-nm red laser.

### 3.5 Enzyme-linked immunosorbent assay (ELISA) measurement of relative antibody titre

We designed an indirect ELISA and calculated the relative antibody titre from each fish plasma sample according to a method established by Sacks et al. (1988).^50^ Our objective was to quantify the amount of *S. molnari*-specific IgM in fish plasma and hence the antigen coating remained constant throughout the experiment. We collected and pooled together parasite lysate derived from three different individual fish at a 1:1:1 mass ratio. Live parasite BSs were pelleted and resuspended in phosphate-buffered saline (PBS) supplemented with the cOmplete Protease Inhibitor Cocktail (Roche Diagnostics GmbH, Mannheim, Germany) by adding one part (volume-wise) of the manufacturer’s recommended stock solution to 24 parts of PBS. This suspension was subjected to a minimum of three freeze-thaw cycles before the lysate was centrifuged at 13,000 *g* for 10 min at 4 °C. We collected the supernatant and measured the absorbance at 280 nm to estimate the protein content using a NanoDrop 2000 (Thermo Fisher Scientific, Wilmington, DE, USA). This exact antigen pool was aliquoted and frozen until all plasma samples were collected for the ELISA.

The *S. molnari* antigen was diluted to 25 μg/mL in 29 mM Na_₂_CO_₃_ 71 mM NaHCO_₃_ pH 9.6 coating buffer. 100 μL were added to wells of a 96-well flat-bottom immunoGrade plates (Carl Roth GmbH, Karlsruhe, Germany) and left to incubate overnight at 4 °C. The coating solution was decanted the next day and 100 μL of PBS 5% non-fat dry milk powder (5 g/100 mL) was added to each well. We left the plate blocking at ambient temperature for 3 hours. We washed away the blocking buffer three times using 100 μL of PBS 0.1% Tween-20. We then added either 50 μL of PBS (for background subtraction) or experimental fish plasma diluted in PBS in triplicate. The specific details of fish plasma dilution and which samples to include on each plate are explained by Sacks et al. (1988).^50^ Briefly, a dilution series of a seropositive fish plasma from another experiment served as a standard and was included on every plate.^51^ We measured the specific antibody titre of this specimen in an independent experiment through a two-fold plasma dilution series. The last dilution with an absorbance above the mean absorbance plus two SDs of a seronegative SPF naïve cohort (n = 5, plasma diluted 1:100) was considered the antibody titre of the standard. We ensured that we had enough of this plasma for every plate we intended to run onwards. Thus, a two-fold dilution series of the standard, ranging from 1:200 to 1:12,800, was included on every plate for the purpose of determining the relative antibody titre and to account for inter-assay and -plate variation. The experimental fish plasma was generally diluted either 1:100 for the naïve group or 1:200 for the infected group, provided the absorbance at 450 nm (A450nm) fell within the range of the standard curve. Otherwise, a specimen was assayed again at a higher or lower dilution as appropriate. The plasma was incubated for 1 hour at ambient temperature. It was washed away thrice with PBS 0.1% Tween-20. We repeated these incubation steps sequentially with a 100 μL of secondary mouse anti-carp IgM antibody (clone WCI12 supernatant diluted 1:2000 in PBS) and a tertiary anti-mouse IgG-horse radish peroxidase conjugate antibody diluted 1:5000 (Sigma-Aldrich, Darmstadt, Germany) before a final wash step (five washes) and signal development by adding 100 μL of pre-warmed TMB per well (Thermo Fisher Scientific, Rockford, IL, USA). The plates were incubated for 6 min before we stopped the reaction with 2 M sulfuric acid.

We measured the A450nm on the Infinite 200 PRO microplate reader (Tecan, Männedorf, Switzerland). Before analysis, all measurements were background-corrected by subtracting the mean of the triplicate PBS internal negative control measurements included on every plate.

### 3.6 qPCR quantification of parasitemia

We extracted total DNA from 2 μL of whole blood. The blood was immediately stored in 400 μL of TNES-urea (10 mM Tris-HCl, 125 mM NaCl, 10 mM EDTA, 0.5% SDS, and 4 M urea, pH 8.0) at 4 °C. Once the entirety of samples for the experiment had been collected, the DNA was purified via a modified phenol-chloroform extraction as described by Holzer et al. (2004).^52^ We measured the yield and the purity via spectrophotometry (ratio of absorbance at 260 nm to 280 nm) using the NanoDrop 2000 (Thermo Fisher Scientific, Wilmington, DE, USA). All samples were diluted to 100 ng/μL to normalize the input material for the qPCR. We measured relative parasitemia as the quantity of *S. molnari* SSU rDNA relative to *C. carpio* β-actin DNA using the primers and probes in a duplex TaqMan real-time PCR assay as described previously.^24^ Naturally, severe disease would both increase parasitemia and deplete host erythrocytes,^30^ increasing the parasite signal while decreasing the host signal. For relative quantification, specimens in which no parasite DNA was amplified were assigned a Ct value of 50, the number of PCR cycles we programmed. Measurements were made in technical duplicates. The TaqMan qPCR was performed on the QuantStudio 6 (Applied Biosystems, Foster City, CA, USA).

### 3.7 EdU pulse labeling of proliferating and memory B cells

Experimental fish were injected with the thymidine analogue EdU to measure cells proliferating *in vivo* during the acute response to *S. molnari*. The fish in these two groups were each injected intraperitoneally with 100 μL of 5 mM EdU (diluted in a 1:1 sterile solution of distilled H_2_O and RPMI 1640) one day before sampling. We sampled four infected and three mock-infected fish per week at one-week intervals between weeks 1 and 9 post-infection.

A separate group of fish was instead injected three times (at two-day intervals) during a single labeling window/week between weeks 3 and 8 of this experiment. The purpose was to capture the long-term outcome of the proliferation and EdU incorporation events that had occurred during a particular labeling window between different organs and between different windows. Therefore, we sampled these fish approximately six months after each respective labeling window.

The entirety of the head kidney and spleen were biopsied. We placed the tissues upon the surfaces of 100-μm cell strainers (Corning, Durham, NC, USA). The tissues were then gently dissociated using the textured end of a syringe plunger with simultaneous washing and rinsing with RPMI 1640. The cell suspensions were then loaded onto 25% Percoll (Cytiva, Uppsala, Sweden), prepared with one part Percoll and three parts RPMI 1640. The heparinized blood was loaded onto Ficoll-Paque (Cytiva, Uppsala, Sweden). Leukocytes were enriched by density centrifugation at 500 *g* for 20 min at 4 °C with minimum acceleration and braking. We collected either the pellet for splenocytes and head kidney leukocytes, or the buffy coat for blood mononuclear cells. These cell fractions were washed twice in RPMI 1640. We enumerated the cells by Trypan Blue exclusion in a Bürker chamber. One million were allotted per fish per compartment, they were stained for detection of IgM^+^ cells as described for blood leukocytes minus the DiOC6(3) staining.

Cells that had incorporated EdU were detected using the Click-iT EdU Alexa Fluor 488 Flow Cytometry Assay Kit (Life Technologies Limited, Paisley, UK) according to the manufacturer’s instructions except that the entire protocol was conducted on 96-well non-tissue culture-treated polystyrene V-bottom plates (Greiner Bio-One, Les Ulis, France) with the maximum wash volumes adjusted accordingly: we used 10% of the recommended freshly prepared Click-iT cocktail per specimen, and hence resuspended the cells in only 10 µL of saponin-based permeabilization and wash reagent in the step prior to the Click-iT reaction. Specimens were analyzed on the BD FACSCanto II.

### 3.8 Primer selection

For SYBR Green-based qPCR, the *igm*-specific primer pairs (against both *membIgM* and *secIgM*) and housekeeping *bactin*-specific primer pairs were previously described.^24^ We designed the other hypothetical/predicted markers of B cell activation/differentiation and they are described in Table S1.

For the multiplex qPCR, the gene/primer selection was based on a single-cell RNA sequencing (scRNA-seq) B cell atlas published for grass carp *Ctenopharyngodon Idella*.^16^ The nucleotide sequences of the B cell-specific target genes in common carp were derived from the respective NCBI nucleotide accession using the Pyrosequencing assay design software (Biotage AB, Uppsala, Sweden). For each of the 18 target genes, we designed either the sense or the antisense primer to bind an exon*-*exon junction. They are outlined in Table S1. *eef1a1* (eukaryotic translation elongation factor 1 alpha 1) and *rps11* (ribosomal protein S11) were included as reference genes.^53^

### 3.9 Quantitative reverse transcription PCR (RT-qPCR) gene expression profiling of B cell activation and differentiation markers

Fish were dissected and the B cells stained as described for the proliferation assay. MACS was performed according to the manufacturer’s instructions using Anti-Mouse IgG Microbeads (Miltenyi Biotec, Bergisch Gladbach, Germany). Cells were stored in RNAlater (Thermo Fisher Scientific Baltics UAB, Vilnius, Lithuania) overnight, centrifuged the next day at 5,000 *g* either undiluted or diluted 1:1 with PBS. We then stored the pellet at -20 °C for RNA isolation using the NucleoSpin RNA kit (MACHEREY-NAGEL, Düren, Germany) according to the manufacturer’s instructions. RNA was aliquoted for long-term storage at -80 °C or to prepare a 4 ng/µL solution from which 2.5 µL was used (10 ng) in the 10 µL reaction format of the Power SYBR Green RNA-to-CT 1-Step Kit (Thermo Fisher Scientific Baltics UAB, Vilnius, Lithuania) as recommended by the manufacturer. Enough of an equally proportioned pool of 12 randomly selected RNA specimens was prepared and assayed as described above as an inter-run calibrator. We made technical duplicate measurements on the QuantStudio 6 (Applied Biosystems, Foster City, CA, USA). We repeated measurements for any technical duplicates that varied over 0.5 cycles. In this case, Ct values were adjusted based on the inter-run calibrator but otherwise, we followed the sample-maximization method for plate/assay design. We analyzed the data via the 2^−ΔΔCT^ method: all data were initially normalized to the Ct values of *bactin* from the same RNA specimen (ΔCt), these data were further normalized to the mean of the ΔCt of the corresponding control group (ΔΔCt).

### 3.10 Multiplex qPCR gene expression profiling

RNA specimens extracted for one-step RT-qPCR described above were also aliquoted and reverse-transcribed into cDNA using the Reverse Transcription Master Mix (Standard BioTools, San Francisco, CA, USA) in a Thermocycler-T1 (Biometra, Analytik Jena, Jena, Germany) instrument. The resulting cDNA was pre-amplifed using the Preamp Master Mix (Standard BioTools) and a primer master mix with a final concentration of 100 µM per primer pair. Finally, preamplified cDNA was treated with exonuclease I (New England BioLabs, Ipswich, MA, USA) and diluted in SsoFast EvaGreen Supermix with Low ROX (Bio-Rad) and 192.24 DELTAgene Sample Reagent (Standard BioTools).

We used the BioMark HD system (Standard BioTools) to profile the expression of selected B cell marker genes and two reference genes (Table S1) in the preamplified cDNA samples. To this end, the primers and pre-amplified cDNAs were inserted to the assay and sample inlets, respectively, on a 192.24 Gene Expression biochip (Standard BioTools). The Control Line Fluid and the Actuation and Pressure Fluid (Standard BioTools) were transferred to the wells provided on the biochip. Having underwent the Load Mix script using the IFC Controller RX (Standard BioTools), the 192.24 Gene-Expression biochip was finally transferred to the BioMark HD-System (Standard BioTools) to perform the quantification reactions.

The Ct values were retrieved using the Fluidigm RealTime PCR Analysis Software v4.5.2 and translated into copy numbers based on external amplicon-specific standard curves (*R*^2^ > 0.99). The relative copy numbers were normalised by the geometric mean of the expression values of the suitable reference genes *eef1a1* and *rps11*.^53^

### 3.11 Fixation, permeabilization, and intracellular anti-IgM staining of putative plasma cells

For detection of antibody-secreting cells, we used head kidney biopsies from non-IS fish that received either 50,000 or 250,000 *S. molnari* BSs. Tissue processing and staining for membrane-bound/cell surface IgM was as described above but we started with five million cells in anticipation of detecting rare cell populations.

To costain for cell surface and intracellular IgM, we used the same anti-carp IgM clone WCI12 antibody specially purified from supernatant with the Mouse TCS Antibody Purification kit (Abcam, Cambridge, UK) according to the manufacturer’s instructions. The antibody was then biotinylated using EZ-Link Sulfo-NHS-LC-Biotin (Thermo Fisher Scientific, Rockford, IL, USA) according to the manufacturer’s instructions with a 100-fold molar excess of biotin. The prepared antibody (WCI12-biotin) was then dialyzed against a 1000-fold volume excess of PBS. The successful biotinylation reaction was confirmed by flow cytometry and western blot using streptavidin reagents (data not shown).

After cell surface staining, we washed the cells twice with Hanks’ Balanced Salt Solution (HBSS) (Thermo Fisher Scientific, Paisley, UK) supplemented with 1% fetal bovine serum. We then fixed the cells in 100 µL of IC Fixation Buffer (Thermo Fisher Scientific, Carlsbad, CA, USA) for 30 min at ambient temperature. From this point onwards, we pelleted cells by centrifugation at 800 *g* for 5 min. The cells were then washed twice with HBSS 1% fetal bovine serum before permeabilization for 15 min at ambient temperature with Permeabilization Buffer (Thermo Fisher Scientific, Carlsbad, CA, USA). After this fixation and permeabilization, we pelleted cells and incubated them for 30 min in Permeabilization Buffer with 5% heat-inactivated BALB/C mouse serum to saturate the existing anti-mouse IgG reagent used for membrane IgM staining. The cells were pelleted and stained in the refrigerator overnight with 300 ng of WCI12-biotin per specimen.

We washed cells twice with Permeabilization Buffer. Cells were then stained with 120 µL of streptavidin PE-Cyanine7 conjugate (Thermo Fisher Scientific, Carlsbad, CA, USA) diluted 1:500 in Permeabilization Buffer for 30 min in ice. The excess streptavidin reagent was washed away twice in Permeabilization Buffer before we resuspended cells for flow cytometry analysis.

### 3.12 Quantification and statistical analysis

All statistical details of experiments can be found in either the figure legends, or in the supplementary figures and their respective legends.

Throughout this study, statistical significance was always defined by a P value of less than 0.05 or a less than 5% probability of the null hypothesis being true. We used these abbreviations to further specify the outcomes of all statistical tests we performed: ns (not significant); * p < 0.05; ** p < 0.01; *** p < 0.001; **** p < 0.0001.

## Acknowledgments

We would like to acknowledge: the Ministry of Education, Youth and Sports-Interaction, USA of the Czech Republic (MŠMT-LTAUSA) for funding this work through the project LTAUSA19108 granted to T.K.; the Czech Science Foundation (GAČR) for funding this work through the projects 23-08042K and 19-28399X granted to T.K. and A.S.H., respectively. We would also like to acknowledge Mrs. Joana Pimentel for the care, effort, and attention she dedicated to fish husbandry and to this study.

## Authorship contributions

Conceptualization, J.T.H.C., A.P.S., A.R., A.S.H. and T.K.; Methodology, J.T.H.C., A.P.S., A.R., A.S.H. and T.K.; Formal analysis, J.T.H.C. and A.R.; Investigation, J.T.H.C., A.P.S., N.D., J.M., A.R. and T.K.; Data curation, A.R.; Writing – Original Draft, J.T.H.C.; Writing – Review & Editing, J.T.H.C., A.P.S., N.D., J.M., A.R., A.S.H. and T.K.; Visualization, J.T.H.C. and A.R.; Supervision, A.R., A.S.H. and T.K.; Project administration, A.S.H. and T.K.; Funding Acquisition, A.S.H. and T.K.

## References

1. Flajnik, M.F. (2018). A cold-blooded view of adaptive immunity. Nat Rev Immunol 18, 438–453. 10.1038/s41577-018-0003-9.

2. Pradeu, T., and Du Pasquier, L. (2018). Immunological memory: What’s in a name? Immunol Rev 283, 7–20. 10.1111/imr.12652.

3. Perey, D.Y., Finstad, J., Pollara, B., and Good, R.A. (1968). Evolution of the immune response. VI. First and second set skin homograft rejections in primitive fishes. Lab Invest 19, 591–597.

4. Linthicum, D.S., and Hildemann, W.H. (1970). Immunologic responses of Pacific hagfish. 3. Serum antibodies to cellular antigens. J Immunol 105, 912–918.

5. Sigel, M.M., Voss, E.W.J., and Rudikoff, S. (1972). Binding properties of shark immunoglobulins. Comp Biochem Physiol A Comp Physiol 42, 249–259. 10.1016/0300-9629(72)90384-2.

6. Eve, O., Matz, H., and Dooley, H. (2020). Proof of long-term immunological memory in cartilaginous fishes. Dev Comp Immunol 108, 103674. 10.1016/j.dci.2020.103674.

7. Dooley, H., and Flajnik, M.F. (2005). Shark immunity bites back: affinity maturation and memory response in the nurse shark, Ginglymostoma cirratum. Eur J Immunol 35, 936–945. 10.1002/eji.200425760.

8. Schaperclaus, W. (1942). Beitrag zur Kenntnis der Punctata-Formen und -Typen und zur Theorie der Entstehung der infektiosen Bauchwassersucht des Karpfens VII Untersuchungen uber die ansteckende Bauchwassersucht des Karpfens und ihre Bekampfung. Zentralblatt fuer Bakteriologie Jena Abt II 105, 49–72.

9. Śnieszko, S. Badania bakteriologiczne i serologiczne nad bakteriami posocznicy karpi, 1938;.

10. Van Muiswinkel, W.B. (2008). A history of fish immunology and vaccination I. The early days. Fish Shellfish Immunol 25, 397–408. 10.1016/j.fsi.2008.02.019.

11. Davidson, G.A., Lin, S.H., Secombes, C.J., and Ellis, A.E. (1997). Detection of specific and ’constitutive’ antibody secreting cells in the gills, head kidney and peripheral blood leucocytes of dab (Limanda limanda). Vet Immunol Immunopathol 58, 363–374. 10.1016/s0165-2427(97)00017-2.

12. Ye, J., Kaattari, I.M., and Kaattari, S.L. (2011). The differential dynamics of antibody subpopulation expression during affinity maturation in a teleost. Fish Shellfish Immunol 30, 372–377. 10.1016/j.fsi.2010.11.013.

13. Ma, C., Ye, J., and Kaattari, S.L. (2013). Differential compartmentalization of memory B cells versus plasma cells in salmonid fish. Eur J Immunol 43, 360–370. 10.1002/eji.201242570.

14. Bromage, E.S., Kaattari, I.M., Zwollo, P., and Kaattari, S.L. (2004). Plasmablast and plasma cell production and distribution in trout immune tissues. J Immunol 173, 7317–7323. 10.4049/jimmunol.173.12.7317.

15. Wu, L., Fu, S., Yin, X., Guo, Z., Wang, A., and Ye, J. (2019). Long-Lived Plasma Cells Secrete High-Affinity Antibodies Responding to a T-Dependent Immunization in a Teleost Fish. Front Immunol 10, 2324. 10.3389/fimmu.2019.02324.

16. Pan, Y., Wu, C., Zhong, Y., Zhang, Y., and Zhang, X. (2023). An Atlas of Grass Carp IgM+ B Cells in Homeostasis and Bacterial Infection Helps to Reveal the Unique Heterogeneity of B Cells in Early Vertebrates. J Immunol 211, 964–980. 10.4049/jimmunol.2300052.

17. Born-Torrijos, A., Kosakyan, A., Patra, S., Pimentel-Santos, J., Panicucci, B., Chan, J.T.H., Korytar, T., and Holzer, A.S. (2022). Method for Isolation of Myxozoan Proliferative Stages from Fish at High Yield and Purity: An Essential Prerequisite for In Vitro, In Vivo and Genomics-Based Research Developments. Cells 11, 377. doi: 10.3390/cells11030377. 10.3390/cells11030377.

18. Dobai, T., and Bartosova-Sojkova, P. (2024). Sphaerospora molnari. Trends Parasitol 40, 352–353. 10.1016/j.pt.2023.12.011.

19. Siddall, M.E., Martin, D.S., Bridge, D., Desser, S.S., and Cone, D.K. (1995). The demise of a phylum of protists: phylogeny of Myxozoa and other parasitic cnidaria. J Parasitol 81, 961–967.

20. Holland, J.W., Okamura, B., Hartikainen, H., and Secombes, C.J. (2011). A novel minicollagen gene links cnidarians and myxozoans. Proc Biol Sci 278, 546–553. 10.1098/rspb.2010.1301.

21. Foox, J., and Siddall, M.E. (2015). The Road To Cnidaria: History of Phylogeny of the Myxozoa. J Parasitol 101, 269–274. 10.1645/14-671.1.

22. Yahalomi, D., Atkinson, S.D., Neuhof, M., Chang, E.S., Philippe, H., Cartwright, P., Bartholomew, J.L., and Huchon, D. (2020). A cnidarian parasite of salmon (Myxozoa: Henneguya) lacks a mitochondrial genome. Proc Natl Acad Sci U S A 117, 5358–5363. 10.1073/pnas.1909907117.

23. Abos, B., Estensoro, I., Perdiguero, P., Faber, M., Hu, Y., Diaz Rosales, P., Granja, A.G., Secombes, C.J., Holland, J.W., and Tafalla, C. (2018). Dysregulation of B Cell Activity During Proliferative Kidney Disease in Rainbow Trout. Front Immunol 9, 1203. 10.3389/fimmu.2018.01203.

24. Korytar, T., Wiegertjes, G.F., Zuskova, E., Tomanova, A., Lisnerova, M., Patra, S., Sieranski, V., Sima, R., Born-Torrijos, A., Wentzel, A.S., et al. (2019). The kinetics of cellular and humoral immune responses of common carp to presporogonic development of the myxozoan Sphaerospora molnari. Parasit Vectors 12, 208–3. 10.1186/s13071-019-3462-3.

25. Taggart-Murphy, L., Alama-Bermejo, G., Dolan, B., Takizawa, F., and Bartholomew, J. (2021). Differences in inflammatory responses of rainbow trout infected by two genotypes of the myxozoan parasite Ceratonova shasta. Dev Comp Immunol 114, 103829. 10.1016/j.dci.2020.103829.

26. Perez-Cordon, G., Estensoro, I., Benedito-Palos, L., Calduch-Giner, J.A., Sitja-Bobadilla, A., and Perez-Sanchez, J. (2014). Interleukin gene expression is strongly modulated at the local level in a fish-parasite model. Fish Shellfish Immunol 37, 201–208. 10.1016/j.fsi.2014.01.022.

27. Abd-Elfattah, A., Kumar, G., Soliman, H., and El-Matbouli, M. (2014). Persistence of Tetracapsuloides bryosalmonae (Myxozoa) in chronically infected brown trout Salmo trutta. Dis Aquat Organ 111, 41–49. 10.3354/dao02768.

28. Houghton, G., and Matthews, R.A. (1990). Immunosuppression in juvenile carp, Cyprinus carpio L.: the effects of the corticosteroids triamcinolone acetonide and hydrocortisone 21-hemisuccinate (cortisol) on acquired immunity and the humoral antibody response to Ichthyophthirius multifiliis Fouquet. J Fish Dis 13, 269–280. 10.1111/j.1365-2761.1990.tb00783.x.

29. Houghton, G., and Matthews, R.A. (1986). Immunosuppression of carp (Cyprinus carpio L.) to ichthyophthiriasis using the corticosteroid triamcinolone acetonide. Vet Immunol Immunopathol 12, 413–419. 0165-2427(86)90148-0.

30. Korytar, T., Chan, J.T.H., Vancova, M., and Holzer, A.S. (2019). Blood feast: Exploring the erythrocyte-feeding behaviour of the myxozoan Sphaerospora molnari. Parasite Immunol 10.1111/pim.12683.

31. Shibasaki, Y., Afanasyev, S., Fernandez-Montero, A., Ding, Y., Watanabe, S., Takizawa, F., Lamas, J., Fontenla-Iglesias, F., Leiro, J.M., Krasnov, A., et al. (2023). Cold-blooded vertebrates evolved organized germinal center-like structures. Sci Immunol 8, eadf1627. 10.1126/sciimmunol.adf1627.

32. Picard-Sanchez, A., Estensoro, I., Del Pozo, R., Piazzon, M.C., Palenzuela, O., and Sitja-Bobadilla, A. (2019). Acquired protective immune response in a fish-myxozoan model encompasses specific antibodies and inflammation resolution. Fish Shellfish Immunol 90, 349–362. 10.1016/j.fsi.2019.04.300.

33. Zwollo, P. (2011). Dissecting teleost B cell differentiation using transcription factors. Dev Comp Immunol 35, 898–905. 10.1016/j.dci.2011.01.009.

34. Zwollo, P., Mott, K., and Barr, M. (2010). Comparative analyses of B cell populations in trout kidney and mouse bone marrow: establishing “B cell signatures“. Dev Comp Immunol 34, 1291–1299. 10.1016/j.dci.2010.08.003.

35. Zwollo, P., Haines, A., Rosato, P., and Gumulak-Smith, J. (2008). Molecular and cellular analysis of B-cell populations in the rainbow trout using Pax5 and immunoglobulin markers. Dev Comp Immunol 32, 1482–1496. 10.1016/j.dci.2008.06.008.

36. Barr, M., Mott, K., and Zwollo, P. (2011). Defining terminally differentiating B cell populations in rainbow trout immune tissues using the transcription factor XbpI. Fish Shellfish Immunol 31, 727–735. 10.1016/j.fsi.2011.06.018.

37. Tafalla, C., Gonzalez, L., Castro, R., and Granja, A.G. (2017). B Cell-Activating Factor Regulates Different Aspects of B Cell Functionality and Is Produced by a Subset of Splenic B Cells in Teleost Fish. Front Immunol 8, 295. 10.3389/fimmu.2017.00295.

38. Granja, A.G., Holland, J.W., Pignatelli, J., Secombes, C.J., and Tafalla, C. (2017). Characterization of BAFF and APRIL subfamily receptors in rainbow trout (Oncorhynchus mykiss). Potential role of the BAFF / APRIL axis in the pathogenesis of proliferative kidney disease. PLoS One 12, e0174249. 10.1371/journal.pone.0174249.

39. Chakravarti, R., and Adams, J.C. (2006). Comparative genomics of the syndecans defines an ancestral genomic context associated with matrilins in vertebrates. BMC Genomics 7, 83–83. 10.1186/1471-2164-7-83.

40. Zwollo, P. (2012). Why spawning salmon return to their natal stream: the immunological imprinting hypothesis. Dev Comp Immunol 38, 27–29. 10.1016/j.dci.2012.03.011.

41. Igarashi, H., Medina, K.L., Yokota, T., Rossi, M.I.D., Sakaguchi, N., Comp, P.C., and Kincade, P.W. (2005). Early lymphoid progenitors in mouse and man are highly sensitive to glucocorticoids. Int Immunol 17, 501–511. 10.1093/intimm/dxh230.

42. Akkaya, M., Kwak, K., and Pierce, S.K. (2019). B cell memory: building two walls of protection against pathogens. Nat Rev Immunol 10.1038/s41577-019-0244-2.

43. Ye, J., Kaattari, I., and Kaattari, S. (2011). Plasmablasts and plasma cells: reconsidering teleost immune system organization. Dev Comp Immunol 35, 1273–1281. 10.1016/j.dci.2011.03.005.

44. Yu, X., Tsibane, T., McGraw, P.A., House, F.S., Keefer, C.J., Hicar, M.D., Tumpey, T.M., Pappas, C., Perrone, L.A., Martinez, O., et al. (2008). Neutralizing antibodies derived from the B cells of 1918 influenza pandemic survivors. Nature 455, 532–536. 10.1038/nature07231.

45. Weisel, F.J., Zuccarino-Catania, G.V., Chikina, M., and Shlomchik, M.J. (2016). A Temporal Switch in the Germinal Center Determines Differential Output of Memory B and Plasma Cells. Immunity 44, 116–130. S1074-7613(15)00505-1 [pii].

46. Ye, J., Bromage, E.S., and Kaattari, S.L. (2010). The strength of B cell interaction with antigen determines the degree of IgM polymerization. J Immunol 184, 844–850. 10.4049/jimmunol.0902364.

47. Vijay, R., Guthmiller, J.J., Sturtz, A.J., Surette, F.A., Rogers, K.J., Sompallae, R.R., Li, F., Pope, R.L., Chan, J.A., de Labastida Rivera, F., et al. (2020). Infection-induced plasmablasts are a nutrient sink that impairs humoral immunity to malaria. Nat Immunol 21, 790–801. 10.1038/s41590-020-0678-5.

48. Holzer, A.S., Piazzon, M.C., Barrett, D., Bartholomew, J.L., and Sitja-Bobadilla, A. (2021). To React or Not to React: The Dilemma of Fish Immune Systems Facing Myxozoan Infections. Front Immunol 12, 734238. 10.3389/fimmu.2021.734238.

49. Secombes, C.J., van Groningen, J.J., and Egberts, E. (1983). Separation of lymphocyte subpopulations in carp Cyprinus carpio L. by monoclonal antibodies: immunohistochemical studies. Immunology 48, 165–175.

50. Sacks, J.M., Gillette, K.G., and Frank, G.H. (1988). Development and evaluation of an enzyme-linked immunosorbent assay for bovine antibody to Pasteurella haemolytica: constructing an enzyme-linked immunosorbent assay titer. Am J Vet Res 49, 38–41.

51. Ganeva, V.O., Korytar, T., Peckova, H., McGurk, C., Mullins, J., Yanes-Roca, C., Gela, D., Lepic, P., Policar, T., and Holzer, A.S. (2020). Natural Feed Additives Modulate Immunity and Mitigate Infection with Sphaerospora molnari (Myxozoa:Cnidaria) in Common Carp: A Pilot Study. Pathogens 9, 1013. 10.3390/pathogens9121013.

52. Holzer, A.S., Sommerville, C., and Wootten, R. (2004). Molecular relationships and phylogeny in a community of myxosporeans and actinosporeans based on their 18S rDNA sequences. Int J Parasitol 34, 1099–1111. 10.1016/j.ijpara.2004.06.002.

53. Adamek, M., Matras, M., Rebl, A., Stachnik, M., Falco, A., Bauer, J., Miebach, A., Teitge, F., Jung-Schroers, V., Abdullah, M., et al. (2022). Don’t Let It Get Under Your Skin! - Vaccination Protects the Skin Barrier of Common Carp From Disruption Caused by Cyprinid Herpesvirus 3. Front Immunol 13, 787021. 10.3389/fimmu.2022.787021.

