## Supplemental Figures for "Immunological memory in a teleost fish: common carp IgM^+^ B cells differentiate into memory and plasma cells"

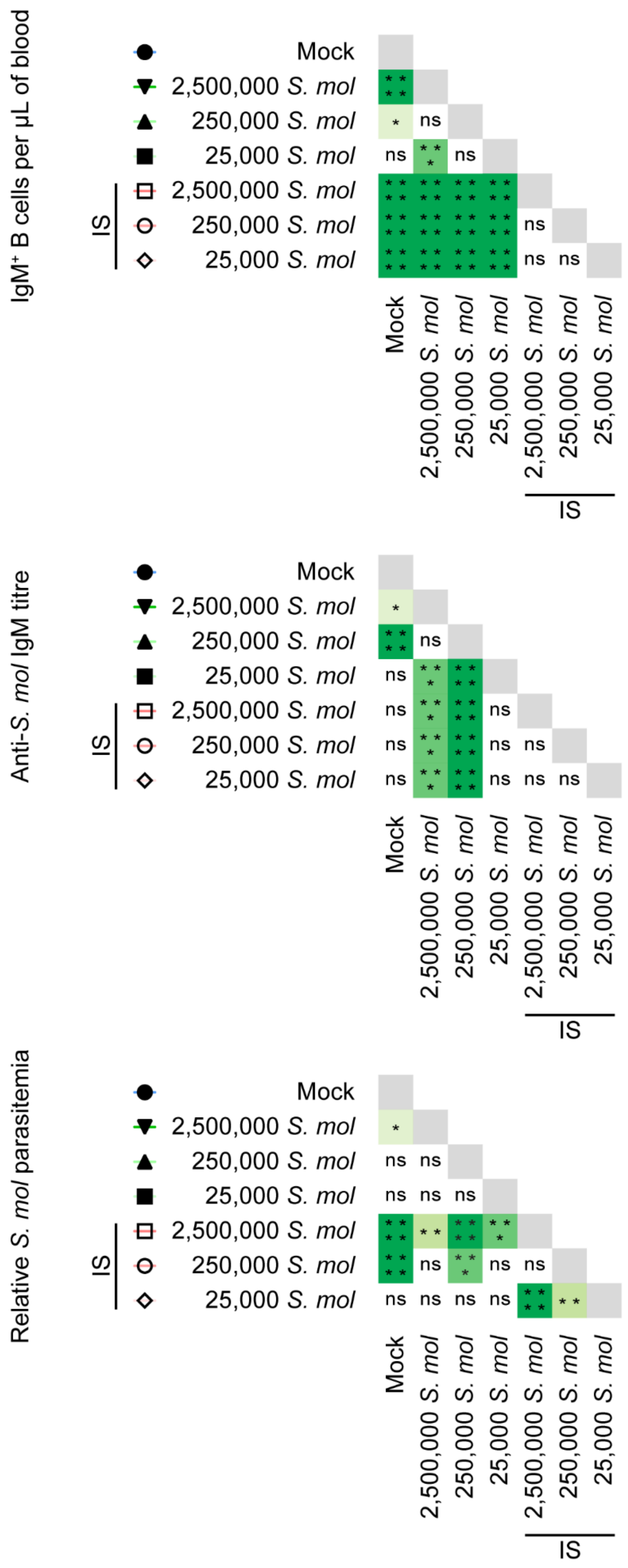


**Figure S1. Statistical analyses of the impact of *S. molnari* (*S. mol*) infection and immunosuppression on blood B cell cellularity of common carp, anti-*S. molnari* IgM antibody titres, and parasitemia.** The analyses presented here correspond to the data presented in the main Figure 2. We performed a two-way ANOVA with a *post hoc* Tukey's multiple comparisons test to make all possible comparisons between infection, the size of the *S. molnari* inoculum and immunosuppression (IS). n ≥ 5 biological replicates per group per timepoint. Gray boxes are comparisons that we did not make because they are between the same groups. ns (not significant); * $p \leq 0.05$; ** $p \leq 0.01$; *** $p \leq 0.001$; **** $p \leq 0.0001$.

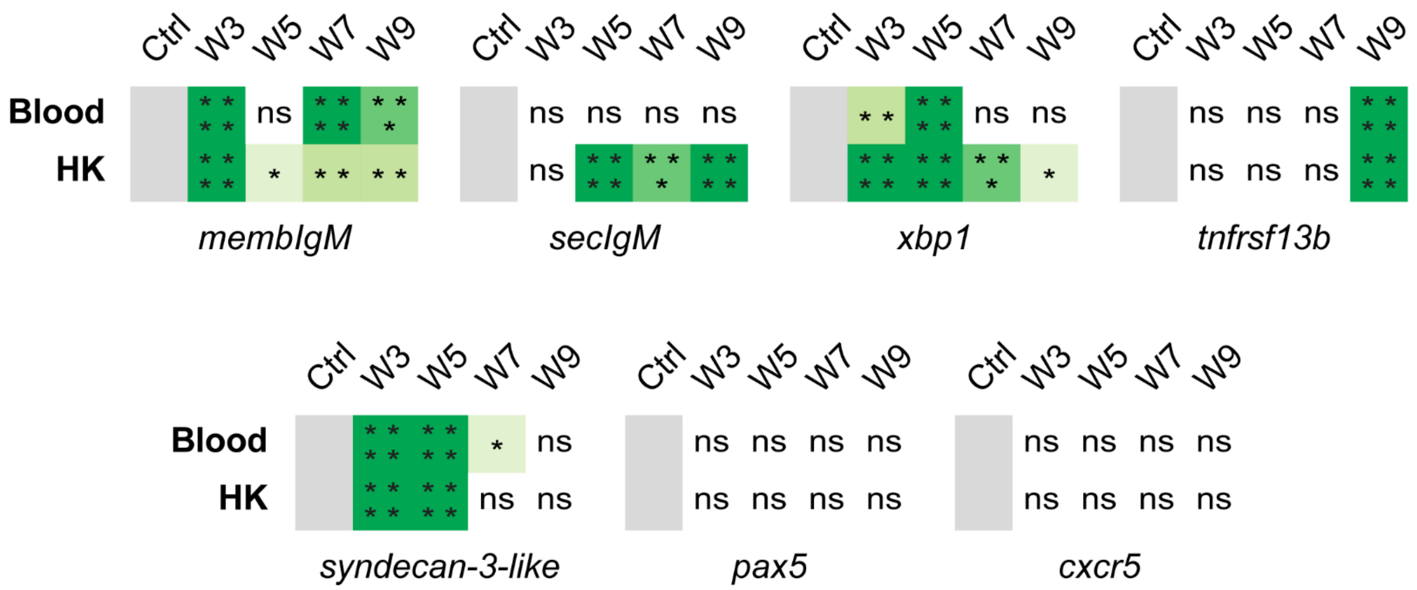


**Figure S2. Statistical analyses of the change in expression of several predicted *Cyprinus carpio* B cell markers of differentiation and survival following *S. molnari* infection.** This figure corresponds to the main Figure 4. A *post hoc* Dunnett's multiple comparisons test compared the relative gene expression ($2^{-\Delta\Delta CT}$) at the indicated timepoints (in weeks) relative to the expression of the same genes in the baseline control group (Ctrl) sampled prior to infection. We refer to and presented relevant two-way ANOVA test results in the Results section. The control group is in gray and was not compared to itself. n ≥ 4 biological replicates per group per timepoint. ns (not significant); * $p \leq 0.05$; ** $p \leq 0.01$; *** $p \leq 0.001$; **** $p \leq 0.0001$.

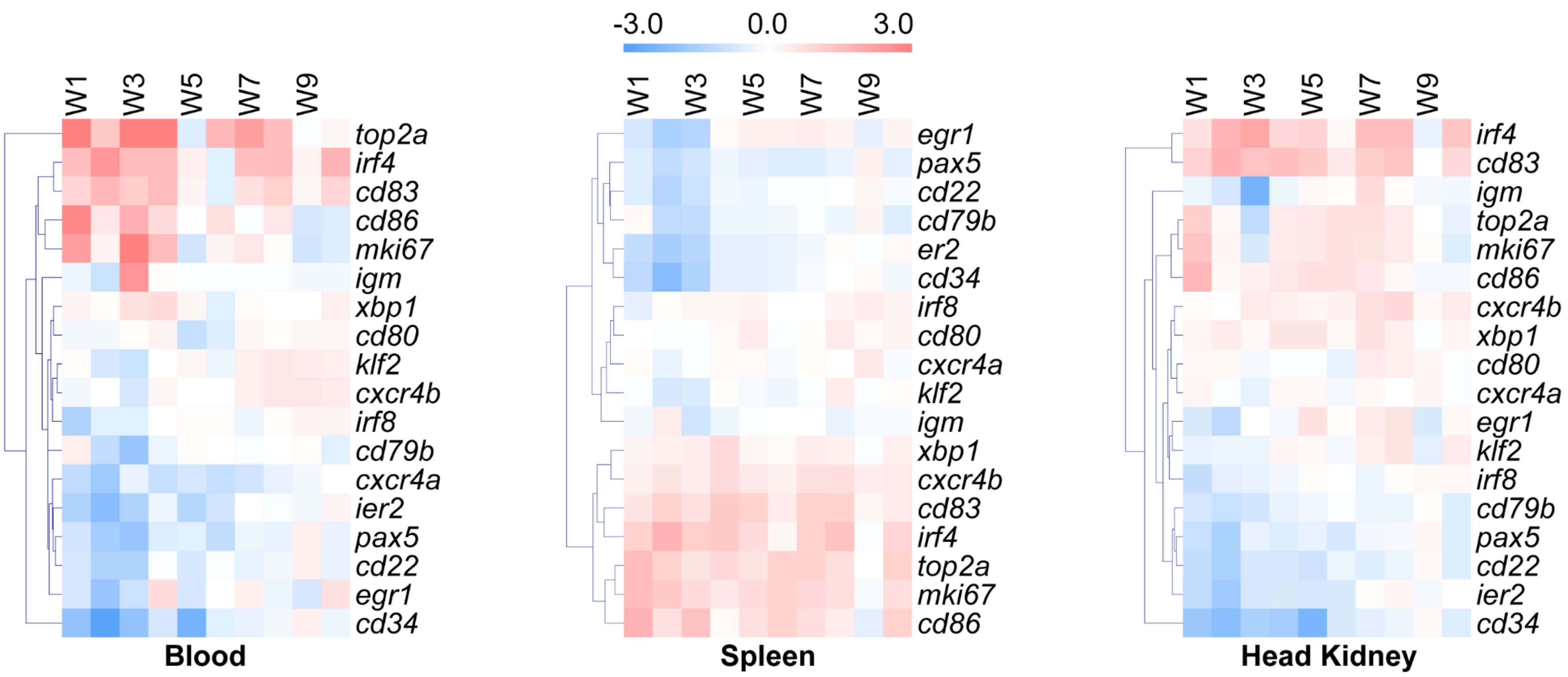


**Figure S3. Hierarchical clustering of the expression of B cell markers in the blood, spleen and head kidney of common carp at various timepoints after *S. molnari* infection.** These *Cyprinus carpio* orthologues were chosen because they were identified by scRNA-seq analysis of the $IgM^+$ B cells of another cyprinid species and define at least six major B cell populations[1]. What we present here is data from the main Figure 5 hierarchically clustered within each compartment by Euclidian distance and an average linkage clustering method with optimization of leaf order (MultiExperiment Viewer 4.9.0). The color code or gradient above represents logarithmically transformed gene copy numbers relative to the control group.

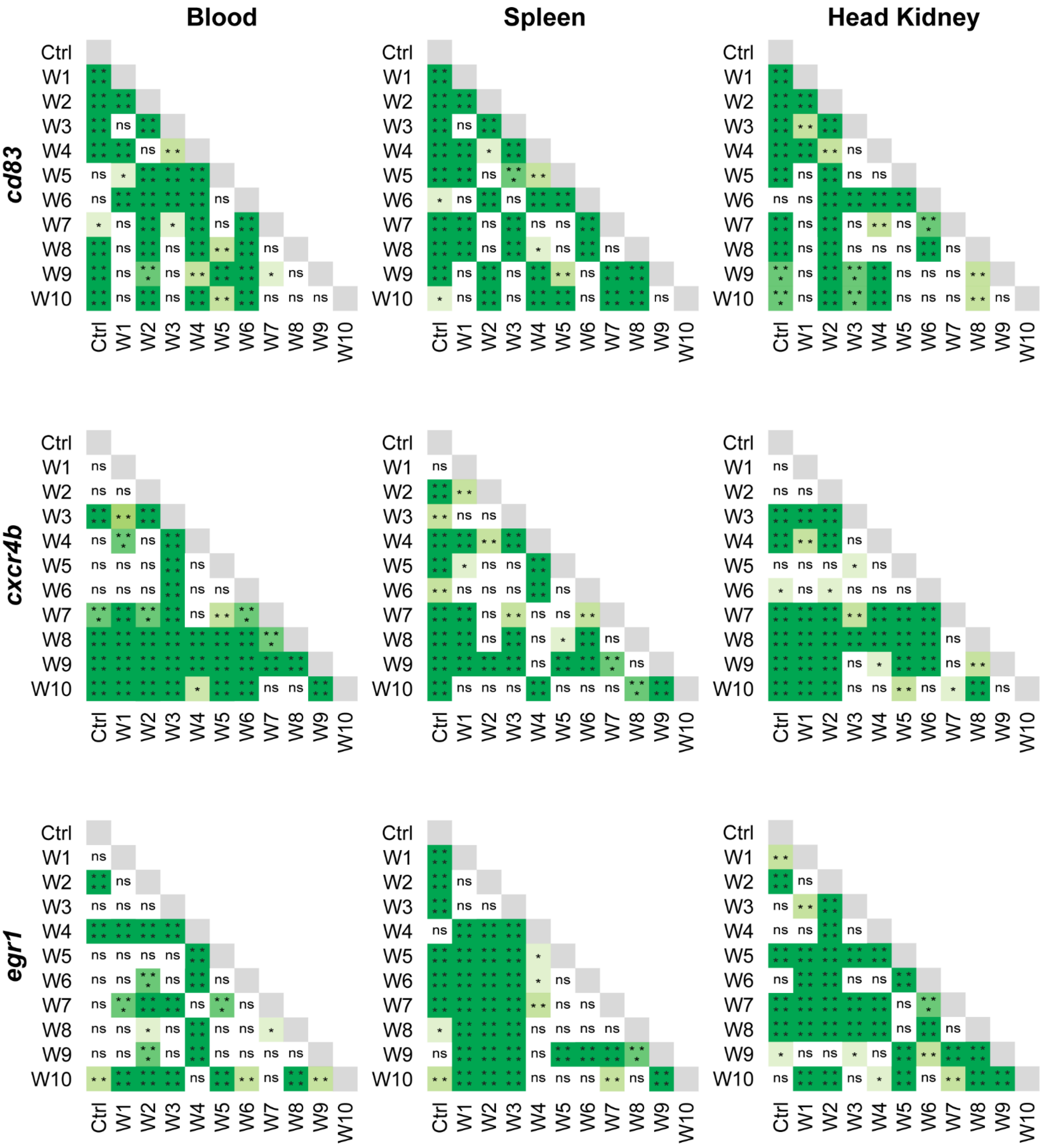


**Figure S4. Statistical analyses of the relative number of copies of several predicted *Cyprinus carpio* B cell markers after *S. molnari* infection in three compartments.** We performed a *post hoc* Holm-Šídák multiple comparisons test after a two-way ANOVA to make every possible comparison between control (Ctrl) and the ten timepoints (in weeks). This is a companion figure to Figure 5. Gray boxes are excluded self-comparisons. n ≥ 4 biological replicates per group per timepoint. ns (not significant); * $p \leq 0.05$; ** $p \leq 0.01$; *** $p \leq 0.001$; **** $p \leq 0.0001$. (continued on next page)

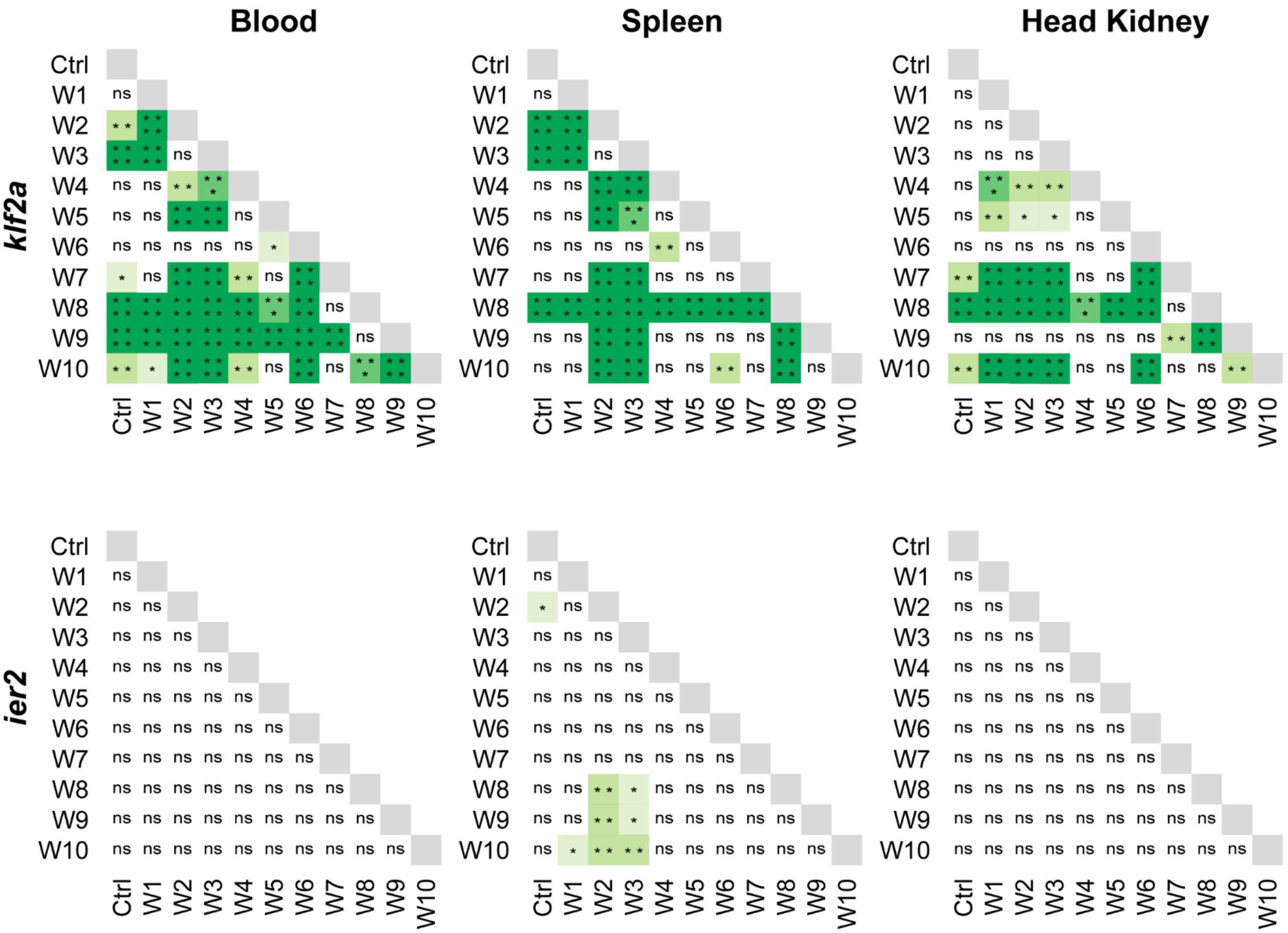


**Figure S4.** (continued)

**Reference**


1. Pan, Y., Wu, C., Zhong, Y., Zhang, Y., and Zhang, X. (2023). An Atlas of Grass Carp IgM+ B Cells in Homeostasis and Bacterial Infection Helps to Reveal the Unique Heterogeneity of B Cells in Early Vertebrates. J Immunol *211*, 964–980. https://doi.org/10.4049/jimmunol.2300052.
