## Supplemental Table for "Immunological memory in a teleost fish: common carp IgM^+^ B cells differentiate into memory and plasma cells"

**Table S1. Summary of all the oligonucleotides designed for this study against various (predicted) B cell or B cell-related markers.** Oligonucleotides are divided into two groups: the first five primers target predicted (in common carp) B cell-associated markers whose expression we measured as part of Figure 4 and the Method Details section ‘Quantitative reverse transcription PCR (RT-qPCR) gene expression profiling of B cell activation and differentiation markers’; the second group includes primers targeting 18 *Cyprinus carpio* orthologues of grass carp B cell-associated genes that Pan et al. (2023)^1^ identified by single-cell RNA sequencing as defining and clustering distinct head kidney IgM^+^ B cell populations. We measured the latter group of genes as part of Figure 5 and the Method Details section ‘Multiplex qPCR gene expression profiling’. These two groups of primers are divided by a dashed line.

| **Gene symbol** | **NCBI or GenBank accession code** | **Gene product** | **Forward (F) and reverse (R) primers (5’-3’)** | **Amplicon length (bp)** |
| --- | --- | --- | --- | --- |
| *tnfrsf13b* | XM_042734762.1 | *Cyprinus carpio* tumor necrosis factor receptor superfamily member 13B | F: CAGTGCTCCGAGCTGTGT  R: AGTACAGCGCTGATCCGGA | 157 |
| *xbp1* | XM_042753507.1 | *Cyprinus carpio* X-box binding protein 1 | F: AAAGCACTTCGAAGGAAACTGAAG  R: CGAGCTCCAACTCCAAGACTT | 109 |
| *syndecan-3-like* | XM_042777052.1 | *Cyprinus carpio* syndecan-3-like | F: GTCTCTTATGGCCACATTTACTCG  R: CCCACAGCAATAATGTCCCAAAA | 122 |
| *pax5* | XM_019109396.2  XM_019109382.2  XM_019109370.2  XM_019109360.2 | *Cyprinus carpio* paired box 5, transcript variants X1, X2, X3, and X4 | F: GGCTATGGCGTCTTTGGCT  R: TCGCGACCAGACACAAGTG | 123 |
| *cxcr5* | XM_042743592.1  XM_042743591.1 | Cyprinus carpio C-X-C chemokine receptor type 5, transcript variants X1 and X2 | F: CTTAAACGGCGGAGGAACCT  R: GCCCAATAGCTTGCAGAGGA | 150 |
| *cd22* | XM_042754392, [XM_042726754](https://www.ncbi.nlm.nih.gov/nucleotide/XM_042726754.1?report=genbank&log$=nuclalign&blast_rank=10&RID=4C3RXRTW016) | *Cyprinus carpio* cluster of differentiation 22 | F: CATTCTCTTGCTGATGTCCTTCAT  R: ATGAGCGTGTGTTCAGAGGAGA | 99 |
| *cd34* | XM_042713470 | *Cyprinus carpio* cluster of differentiation 34 | F: CTTCACAGTTGCTGGGGACAG  R: AACAGGTTCAAAAGCAGGTCCAA | 109 |
| *cd79b* | XM_042767177 | *Cyprinus carpio* cluster of differentiation 79b | CCTCTCAGCTGAAGTAAATATCC,  TGACATCTGATCCATCTTTGACG | 109 |
| *cd80* | XM_042740109 | *Cyprinus carpio* cluster of differentiation 80 | CACAGCTGAGTACAGCGTTCC,  AGGTGGACGGGGCTTAATCAAA | 143 |
| *cd83* | XM_019119505 | *Cyprinus carpio* cluster of differentiation 83 | CCGTCATATGGTACAAGGTTTCT,  TGTCTAATGTAACTGCAGAGGAC | 173 |
| *cd86* | XM_042778525 | *Cyprinus carpio* cluster of differentiation 86 | GTGCGCCATTCTTCATTAAGGG, CTACATGAGGACGTCAGACTAG | 148 |
| *cxcr4a* | XM_042725575 | *Cyprinus carpio* C-X-C chemokine receptor type 4 alpha | TACGAACACATCGTCTTTGAAGAT, GAGCCAACTTTGAGGTTCCGTG | 92 |
| *cxcr4b* | XM_042763477 | *Cyprinus carpio* C-X-C chemokine receptor type 4 beta | CTTCTTCATCTGTTGGTTGCCTTA,  TGCTGTCTCAACCCCATCCTTTA | 165 |
| *egr1* | XM_019064348 | *Cyprinus carpio* early growth response protein1 | ACTGGAGACACGCTTTCAGAAAT,  TCTTACACGGGCCGTTTCACC | 102 |
| *ier2* | XM_019067280 | *Cyprinus carpio* immediate early response protein 2 | CTCTAGAAAGCGACGGAGCAAA,  CGTGCCTATGCCAAGAACAATTG | 173 |
| *ighm* | [AB004106](https://www.ncbi.nlm.nih.gov/nucleotide/AB004106.1?report=genbank&log$=nuclalign&blast_rank=4&RID=4C5G5EWR013), AB004107, MH352354, MH352353 | *Cyprinus carpio* immunoglobulin M heavy chain membrane-bound and secretory protein | CGAATATGCAGTTCCTATTCAAGAT  AACATGTGGAACTTGATGCCCC | 216 |
| *irf4* | XM_019126566 | *Cyprinus carpio* interferon regulatory factor 4 | GGCCCTCTCAGATTACCGCTTA,  TAGTCCAGAGGGCTGTCAGCT | 93 |
| *irf8* | XM_042754085, XM_042743728 | *Cyprinus carpio* interferon regulatory factor 8 | ATCTTCAAAGCGTGGGCGATATT,  ATTTTGAGGAAGTTACTGACCGAT | 130 |
| *klf2* | XM_019065449 | *Cyprinus carpio* Krüppel like factor 2 alpha | GAAAACAGGTGGAAGGAGGAAC, TTATTCTGGCCAACACTGTGGG | 146 |
| *mki67* | XM_042735109 | *Cyprinus carpio* marker of proliferation Ki-67 | GTTCGGAAGGAAGCTGGACTG, AGAACAAGGAGCTCATTTTGACC | 103 |
| *pax5* | XM_019109360 | *Cyprinus carpio* paired box 5 | CAACAGGATCATTCGCACTAAAG, GTGACCCAGGTATCTGCAGTAA | 107 |
| *top2a* | XM_042735154 | *Cyprinus carpio* DNA topoisomerase II alpha | ACTCAGCAAATGTGGGTGTTTGAT, GGTTAACATTGACTCGGAGAATAA | 170 |
| *xbp1* | XM_042753507 | *Cyprinus carpio* X-box binding protein 1 | TTGGAGTTGGAGCTCGAGAATC,  GACTGGGGTTAGATACCCTGG | 121 |

1. Pan, Y., Wu, C., Zhong, Y., Zhang, Y., and Zhang, X. (2023). An Atlas of Grass Carp IgM+ B Cells in Homeostasis and Bacterial Infection Helps to Reveal the Unique Heterogeneity of B Cells in Early Vertebrates. J Immunol *211*, 964–980. <https://doi.org/10.4049/jimmunol.2300052.>
